# Guard cell photorespiration has a major impact on photosynthesis, growth and stomatal behavior in Arabidopsis

**DOI:** 10.1101/2025.01.21.634093

**Authors:** Hu Sun, Nils Schmidt, Tracy Lawson, Martin Hagemann, Stefan Timm

**Affiliations:** University of Rostock, Plant Physiology Department, Albert-Einstein-Straße 3, D-18059 Rostock, Germany; University of Essex, Wivenhoe Park, Colchester CO4 3SQ, UK

**Keywords:** Arabidopsis, environmental acclimation, glycine decarboxylase, guard cells, photosynthesis, photorespiration, stomata

## Abstract

Photorespiration is a mandatory metabolic repair shunt of carbon fixation by the Calvin-Benson (CB) cycle in oxygenic phototrophs. Its extent depends mainly on the CO_2_/O_2_ ratio in chloroplasts, which is regulated via stomatal movements. However, despite comprehensive understanding on the role of photorespiration in mesophyll cells (MC), its role in guard cells (GC) is yet unknown. To analyze this issue, the key enzyme of photorespiration, glycine decarboxylase (GDC), was manipulated through overexpression and antisense suppression of the *GDC* H-protein gene.

A positive correlation of *GDC-H* expression with growth, photosynthesis and carbohydrate biosynthesis was observed in the transgenic lines, demonstrating active photorespiration is involved in stomatal regulation. This view is supported by gas exchange measurements showing that optimized GC photorespiration improves plant acclimation towards conditions requiring a high photorespiratory capacity, including high light. Microscopic analysis revealed that altered photorespiratory flux also affected starch accumulation patterns in GC, eventually serving as the underlying mechanistic for altered stomatal behavior.

Collectively, our data suggest that photorespiration is a key component of the regulatory circuit that coordinates stomatal movements with external and internal CO_2_ availability. Thus, manipulation of photorespiration in GC has the potential to engineer crops maintaining growth and photosynthesis under future climates.

**One-sentence summary:** Guard-cell-specific manipulation of photorespiratory glycine decarboxylase reveals photorespiration is active in guard cells and makes a major contribution to stomatal metabolism and movements.

## Introduction

To enable the biosynthesis of organic compounds, CO_2_ must enter the intracellular air space of leaves and ultimately reach the chloroplasts and site of fixation. This gas exchange is facilitated by stomata, which evolved more than 400 million years ago (Edwards et al., 1992; 1998). Stomata are microscopic, adjustable pores on the leaf surface that regulate CO_2_ influx, while, at the same time, controlling water loss through transpiration (Vavasseur and Raghavendra, 2005; Lawson, 2009; Santelia and Lawson, 2016). Thus, appropriate regulation of stomatal movement is crucial for maintaining plant productivity and water-use efficiency (WUE), especially in response to fluctuations in temperature, light intensity, water availability, and CO_2_ concentration (Pankasem et al., 2024). Stomatal aperture is controlled by two guard cells (GC) surrounding the stomatal pore, which change in turgor and consequently volume to adjust aperture size (Araújo et al., 2011; Lawson and Blatt, 2014). Over the past centuries, significant progress has been made in understanding the development, structure, and physiology of stomata (Daloso et al., 2016; Santelia and Lawson, 2016). Many external, (i.e. atmospheric pressure, light availability and quality, temperature, water availability and humidity, and CO_2_) and internal (i.e. abscisic acid, circadian rhythms, GC ion transport, plant hormones and sugar concentrations) factors affect stomatal behavior (Inoue and Kinoshita, 2017; Jezek and Blatt, 2017; Lawson and Matthews, 2020). However, three main environmental factors regulate stomatal movement during the day. First, illumination typically induces stomatal opening to support photosynthesis. Second, water availability determines the extent to which stomata remain open. Third, CO_2_, often in combination with light, influences stomatal movement over the longer term, with internal CO_2_, *C_i_*, thought to regulate stomatal behavior with mesophyll photosynthesis (Mott, 1988). The mechanisms by which GC sense changes in CO_2_ and translate these signals into adjustments in metabolism is a matter of intense research (Engineer et al., 2016; Takahashi et al., 2022). Although several factors of the signaling cascade have been identified (see below), the actual CO_2_ sensor is still unknown. Additionally, the coordination between GC and mesophyll cell (MC) metabolism and CO_2_ demands is still not fully understood (Santelia and Lawson, 2016).

Stomata open upon illumination in most plants to facilitate CO_2_ uptake for mesophyll photosynthesis. Blue and red light drive this process, with blue light having a dominant, photosynthesis-independent effect at low intensities (∼10 µmol m^-2^ s^-1^). Red light acts at higher intensities, aligning with photosynthesis demands, and is thought to be key in linking MC CO_2_ needs with stomatal behavior, i.e. stomatal conductance (*g_s_*). Interestingly, coordination between photosynthesis and *g_s_* was recently shown to be regulated through *C_i_*, both, by *C_i_*-dependent and *C_i_*-independent mechanisms (Taylor et al., 2024). Chloroplasts regulate red-light responses, but it remains unclear whether this applies specifically to MC or GC chloroplasts (Lawson, 2009). GC photosynthesis, though debated, produces ATP and NADPH via the chloroplast electron transport chain (Lawson et al., 2002; 2003; Lawson, 2009) and supports blue light-induced opening (Santelia and Lawson, 2016). GC chloroplasts also host a functional CB-cycle, yet the extent of GC photosynthetic contribution compared to MC remains uncertain (Lawson, 2009; Lawson et al., 2014). Stomatal closure during water shortage is mediated by abscisic acid (ABA), synthesized within GC using endogenous enzymes. ABA synthesis can be self-stimulated (Munemasa et al., 2015). Light-induced opening and ABA-mediated closure rely on transporter activities for ion movements regulated via distinct signal perception mechanisms: phototropins for blue light and phytochromes or photosynthetic pigments for red light while the ABA-signal is translated via the ABA-signaling pathway (Chen et al., 2012). Stomatal opening occurs through GC turgor increases driven by osmotic active substance accumulation, mainly K⁺, Cl⁻, malate, and sucrose. The proton motive force (PMF), generated by H⁺-ATPase activity, powers ion transport, while sucrose metabolism supplies cytosolic energy and redox equivalents, fueling other GC processes (Roelfsema and Hedrich, 2005; Gaxiola at el., 2007; Daloso et al., 2016). Metabolism of organic acids (malate, fumarate, pyruvate) and carbohydrates (starch, sucrose) in GC is also vital for stomatal dynamics (Araújo et al., 2011; Chen et al., 2012; Li et al., 2014).

In recent years, the significance of GC starch accumulation and its turnover has become a focus of research (Dang et al., 2024). Early observation showed the presence of starch granules correlates with stomatal aperture (Lloyd, 1908), suggesting starch is key to maintain GC movements. Usually, starch is synthesized in plastids of all cell types, including photosynthetically active and heterotrophic, non-photosynthesizing tissues. GC starch metabolism differs from MC, with starch granules being rapidly degraded upon illumination to produce osmotic compounds supporting opening. Recent findings suggest that hydrogen peroxide (H_2_O_2_) is involved in the remobilization of GC starch under optimal conditions, too (Shi et al., 2024; da Silva et al., 2024). Unlike MC, GC retain starch in darkness, crucial for early opening stages. Structural differences between GC and MC chloroplasts (e.g., reduced thylakoid structures) indicate distinct roles (Vavasseur and Raghavendra, 2005; Lawson, 2009; Flütsch et al., 2020). However, experimental evidence highlights the importance of GC photosynthesis in stomatal function, involving CO_2_ fixation via the CB-cycle and phospho*enol*pyruvate carboxylase (PEPC), though the carbon contribution from each pathway remains unclear (Lawson, 2009; Lawson et al., 2014; Daloso et al., 2015).

In addition to light intensity and water availability, stomatal also respond to external CO_2_ concentrations. Experimentally elevated CO_2_ in the atmosphere reduced stomatal aperture, whilst lower CO_2_ opens stomata (Negi et al., 2008; Engineer et al., 2016). GC sensing of CO_2_ and aligning it with MC photosynthesis demand was thought to be based on *C_i_*, however, recent evidence suggests requirement and intense interplay between *Ci*-dependent and *Ci*-independent mechanism, respectively (Taylor et al., 2024). Studies indicate that isolated GC can respond to CO_2_ changes, suggesting they possess the necessary components for CO_2_ perception and signaling (Edwards and Bowling, 1985; Weyers et al., 1983; Hu et al., 2010). The current signaling cascade involves carbonic anhydrases (CA1 and CA4), which mediate high CO_2_-induced stomatal closure through bicarbonate (HCO_3⁻_) as a messenger (Hu et al., 2010, 2015). Other regulatory components include, among others, protein kinases (HT1 - high leaf temperature 1, OST1 - open stoma 1), anion channels (SLAC1 - slow anion channel-associated 1, QUAC1 - quick anion channel 1), and transport proteins (RHC1 MATE - resistant to high carbon dioxide 1 multidrug and toxin extrusion transporter), which affect different stages of the signaling process (Engineer et al., 2016). CO_2_ signaling also intersects with ABA pathways, as suggested by studies on ABA receptor mutants (PYR/RCARs - pyrabactin resistance/regulatory component of ABA receptors) and PP2C protein phosphatases (*abi1-1*, *abi2-1*). However, the nature of this interaction remains enigmatic (Engineer et al., 2016). Ca²⁺ ions may also act as downstream messengers in this regulatory network, supported by the identification of the ABA-insensitive mutant *gca2* (growth controlled by abscisic acid; Young et al., 2006), though the role of *GCA2* requires further investigation.

In plants, the CB-cycle and photorespiration are tightly regulated by the CO_2_/O_2_ ratio (Busch, 2020) in the chloroplasts. Since GC contain RubisCO and are capable of performing photosynthesis (Reckmann et al., 1990; Cardon and Berry, 1992; Lemonnier and Lawson, 2024), functional photorespiration is essential to metabolize 2-phosphoglycolate (2PG), the major byproduct of RubisCO’s oxygenase activity, too. Photorespiration recycles carbon and phosphorus locked in 2PG and prevents its inhibitory effects on key enzymes of carbon utilization such as triosephosphate isomerase (TPI), SBPase, and phosphofructokinase (Kelly and Latzkow, 1976; Flügel et al., 2017; Li et al., 2019). Additionally, photorespiration detoxifies intermediates like glycolate, glyoxylate, and glycine, which otherwise impair RubisCO activation, PSII redox balance, and manganese homeostasis (Timm and Hagemann, 2020). The capacity of photorespiration plays a crucial role in maintaining plant metabolism under environmental fluctuations that affect intracellular CO_2_/O_2_ ratios (Timm et al., 2019; Meacham-Hensold et al., 2024; Sun et al., 2024). To date, the role of photorespiration in GC remained unexplored, whilst investigating this could reveal how GC adapt their metabolism to fluctuating CO_2_/O_2_ ratios and, possibly, align these changes with MC demands. We hypothesize that GC photorespiration eventually contributes to the *C_i_*-dependent regulation mechanism of stomatal movements and communication with the mesophyll. Therefore, we tested if, and how, manipulation of photorespirations central enzyme glycine decarboxylase H-protein (GDC-H) expression in GC has any significant effect on leaf-physiology, metabolism and, particularly, GC movements. Understanding and rationally modify the underlying regulatory circuit could aid in engineering crop plants with higher growth rates and yields under future climates.

## Material and Methods

### Plant material, growth conditions and growth parameters

During this study, *Arabidopsis thaliana* L. (Arabidopsis) ecotype Columbia 0 (Col-0) served as wild-type reference and background to produce guard cell (GC) specific transgenic lines with modulated glycine decarboxylase H-protein (GDC-H) expression. All seeds were surface-sterilized with chloric acid, sown on a soil (Type MiniTray; Einheitserdewerk, Uetersen, Germany) and vermiculite mixture (4:1) and incubated for 2 days at 4°C to break dormancy. Unless stated otherwise, plant growth was performed in controlled environment chambers (Percival) under the following standard conditions: photoperiod - 12/12 h day/night, 22/20°C day/night, ∼120 mmol m^-2^ s^-1^ irradiance, 400 ppm CO_2_ and ∼70% relative humidity. Were specified, the CO_2_ concentration was increased to 3000 ppm or the photoperiod changed (10/14 or 14/10 h day/night), with otherwise equal parameters. During cultivation, plants were watered with 0.2% Wuxal liquid fertilizer (Aglukon) weekly. For all physiological experiment’s plants at growth stage 5.1 (Boyes et al., 2001) were used. The following quantitative growth parameters were determined from 6-10 biological replicates per genotype: **(i)** rosette diameter, longest possible distance of the fully expanded rosette, **(ii)** total leaf-count, only considering fully expanded rosette leaves, and, **(iii)** fresh and dry (dried for approximately 24 h at 100°C) weights of the entire plant rosette.

### Cloning and plant transformation procedures

In order to obtain transgenic lines with GC specific GDC manipulations, we first PCR amplified a 1141 bp genomic fragment of the GC preferential GC1 promotor (Yang et al., 2008) from wildtype DNA using oligonucleotides (sequences in Supp Tab. S5) *At*GC1_S1141_SacI (P950) and *At*GC1-AS-BamHI (P951) and the entire GDC-H coding sequence (489 bp; Kopriva and Bauwe, 1995) from *Flaveria pringlei* cDNA using oligonucleotides *Fp*GDCH-S-PstI (P965) and *Fp*GDCH-AS-PstI (P966). The resulting fragments (*At*GC1-1141 and *Fp*GDCH) were ligated into vector pJET2.1 (Thermo Fisher Scientific) and sequenced for verification. Next, the *At*GC1-1141 SacI-BamHI promoter fragment was excised from pJET2.1:*At*GC1 and ligated into the binary plant transformation vector pGREEN0229, containing the 35S terminator, previously introduced through restriction and ligation from the 35S cassette (EcoRI and EcoRV), to obtain pG0229:*At*GC1:35STer. Finally, the *Fp*GDCH PstI:PstI fragment was excised from pJET2.1:*Fp*GDCH and ligated into pG0229:*At*GC1:35STer in sense (pG0229:AtGC1:*Fp*GDCH:35STer-sense) and antisense (pG0229:*At*GC1:*Fp*GDCH:35STer-antisense) orientation to obtain the GC specific overexpression and antisense suppression constructs (Supp. Fig. S1A, B). Both constructs were introduced into *Agrobacterium tumefaciens* strain GV3101+pSOUP and used for plant transformation (Clough and Bent, 1998). The resulting phosphinotricine (Basta)-resistant plants were PCR verified and stable T3 lines from at least three independent transformation events (designated as GC1:*Fp*GDCH sense lines SL1, SL4 and SL7 and GC1:*Fp*GDCH anti line AL4, AL5 and AL9, respectively) propagated and used for comprehensive characterization.

### Verification of transgenic lines, RT-PCR, and Immunological Studies

To verify the genomic integration of the constructs, leaf DNA, isolated following standard procedures, was PCR amplified (1 min at 94°C, 1 min at 58°C, and 1.0 min at 72°C; 35 cycles) with primers specific for the *Fp*GDCH fragment (P965 for sense and P966 for antisense orientation) in combination with the 35S terminator (P807). The *S16* gene was amplified (1 min at 94°C, 1 min at 58°C, 30 s at 72°C; 35 cycles) using oligonucleotides *S16*-forward (P444) and *S16*-revers (P445) as control (see Supp. Fig. 1C). The functionality of the integrated overexpression and antisense construct was first verified on the whole leaf-basis via semiquantitative RT-PCR, using 2.5 µg leaf RNA for cDNA synthesis (Nucleospin RNA plant kit [Macherey-Nagel] and RevertAid cDNA synthesis kit [MBI Fermentas]). The oligonucleotide combination *Fp*GDCH-S-PstI (P965) and *Fp*GDCH-AS-PstI (P966) was used to amplify the full-length *Fp*GDCH transcript (489-bp PCR product). The constitutively expressed 40S ribosomal protein *S16* gene was amplified with oligonucleotides *S16*-forward (P444) and *S16*-revers (P445) as positive control. Second, changes in GDC-H protein expression was further analyzed by immunoblotting. Briefly, protein extracts of mesophyll cell and GC preparations, in comparison with whole-leaf preparations, were separated by SDS-PAGE and gel blotting experiments performed according to standard protocols. Changes in the protein abundances were detected using specific antibodies for GDC-H (Kopriva et al., 1996), using PGLP1 (Flügel et al., et al., 2017) as a calibration control. ImageJ was used to determine the band intensities as a measure of altered protein abundances from at least 3 independent immunoblots.

### Isolation of mesophyll and guard cell protein extracts

To obtain mesophyll and GC specific protein extracts, we enriched both cell fractions following the protocol described by Lawrence et al. (2019). Briefly, we used 5-week old plants grown under standard conditions. Scotch tape was attached to fully expanded leaved on either, the abaxial (lower, for GC enrichment) and adaxial (upper, for mesophyll enrichment) side of the leaves, respectively. Subsequently, the peels (∼30 per genotype) were gently removed and immediately frozen in liquid nitrogen. For protein extraction, we followed the protocol described earlier (Lawrence et al., 2019). Subsequent to dissolution of the proteins in buffer, their concentration was determined using the BCA Protein Assay Kit (Thermo Scientific) according to the manufacturer’s instructions. For SDS-PAGE and immunoblotting analysis we used 5 µg of the respective protein extracts.

### Standard gas exchange measurements

Photosynthetic gas exchange parameters were determined on a Li-6400xt Portable Photosynthesis System (LI-COR, Lincoln, Nebraska, US), using plants grown under standard conditions. The following settings were used as the standard: 1000 µmol m^−2^ s^−1^ photon flux density (10% blue light), 25°C block temperature, 400 ppm CO_2_, 300 µmol s^−1^ flow rate, and 50–70% relative humidity. To determine CO_2_ compensation points at various O_2_ concentrations (3%, 21%, and 40%; balanced with N_2_), *A*/*C*_i_ curves were measured (400, 300, 200, 100, 50, 25, 0, 400 µL L^−1^ CO_2_). Mean values ± SD of net CO_2_ uptake rates (*A_N_*), CO_2_ compensation points (*Γ*), stomatal conductance (*g*_s_), intercellular CO_2_ concentrations (*C*_i_), transpiration rates (*E*), and, intrinsic water use efficiency (WUE_int_) were consistently calculated from the final 400 µL L^−1^ CO_2_ step from at least 6 biological replicates. Oxygen inhibition of *A_N_* was calculated from measurements at 21% and 40% O_2_ using equation: O_2_ inhibition = (A_21_ – A_40_) / A_21_ × 100. Calculation of *γ* (measure of the photorespiratory CO_2_-release) was performed by linear regression of the *Γ*-versus-oxygen concentration curves and is given as slopes of the respective functions. Light response curves were measured using ambient air CO_2_ and O_2_ levels, using varying light intensities (1759, 1144, 757, 488, 236, 143, 62, 36, 0 µmol m^−2^ s^−1^).

### PSI and PSII chlorophyll fluorescence measurements

Chlorophyll fluorescence measurements were performed on a Dual-PAM 100 (Heinz Walz, Effeltrich, Germany) to determine selected PSI and PSII parameters. We measured chlorophyll fluorescence from the adaxial side of the leaf and it should be noted that PSI refers to P700 measurement on the whole leaf tissue and the PSII is a fluorescence measurement at a certain depth in the leaf tissue. Initial *F*_v_/*F*_m_ (maximum quantum efficiency of PSII) and *P*_m_ (maximum photo-oxidizable P700) values were recorded following a 10 min dark adaptation period. Plants were than exposed to 1000 μmol photons m^−2^ s^−1^ for 10 min to fully induce photosynthesis. Subsequently, light response curves were measured at varying light intensities (1759, 1144, 757, 488, 236, 143, 62, 36, 0 μmol photons m^−2^ s^−1^) at 400 ppm CO_2_ and 21% O_2_.

### Determination of metabolite levels via LC-MS/MS, GC and spectrophotometrically measurements

Abundances of primary metabolites were quantified by liquid chromatography coupled to tandem mass spectrometry (LC-MS/MS) and gas chromatography analysis, using leaf tissue from plants at the end of the day (11 h of illumination). Whole plant-rosettes were harvested under illumination within the growth cabinets, immediately frozen in liquid nitrogen and stored at -80°C. Prior analysis, plants were freeze-dried by lyophilization and aliquoted (∼3-5 mg dry weight). Extraction and LC-MS/MS measurements was carried out using LC-MS grade chemicals as described in Arrivault et al., (2009; 2015) and the modifications specified in Reinholdt et al., (2019). The total amino acid content is a sum parameter of the following representatives: alanine, arginine, asparagine, cysteine, cystine, glutamate, glutamine, glycine, histidine, isoleucine, leucine, lysine, methionine, phenylalanine, proline, serine, threonine, tryptophan, tyrosine, and valine. The total organic acid content is a sum parameter of the following representatives: aconitate, citrate, fumarate, GABA, isocitrate, lactate, malate, and succinate.

Soluble sugars were extracted from homogenized plant material in 800 µl 80% ethanol, containing 20 µg ribitol as internal standard, at 80°C for 30 min. After centrifugation (20,000 g, 10 min, 4°C), the supernatant was dried by lyophilization. The pellet was used for starch quantification through spectrophotometrically analysis using enzymatic assays in ethanolic extracts described elsewhere (Cross et al., 2006). Dried pellets containing soluble carbohydrates were re-suspended in 65 µl pyridine containing 20 mg/ml methoxylamine at 30°C for 90 min. Subsequently, 35 µl N-methyl-N-trimethylsilyl-trifluoracetamide (MSTFA) was added, the samples incubated at 65°C for 90 min and briefly centrifuged (20,000 g, 1 min). After cooling, aliquots of the supernatants were analyzed on the gas-chromatograph 6890N GC System (Agilent technologies), equipped with a TG-5MS column (ThermoScientific). Samples were injected with an inlet temperature of 250°C and 40 ml min^-1^ nitrogen gas flow at a split ratio of 20:1. A flow of 1.8 ml min^-1^ nitrogen gas (mobile phase) and an average velocity of 45 cm s^-1^ at 172 kPa were used. The oven temperature was initially set to 160°C for 2 min, following a gradual increase to 190°C (10°C min^-1^) for 5 min. Subsequently, temperature was increased to 200°C in a step gradient (1°C min^-1^) within 15 min following a gradual increase to 280°C (15°C min^-1^) for 5 min. The Flame Ionization Detector (FID) was set to 250°C, 40 ml min^-1^ hydrogen and 400 ml min^-1^ compressed air and a makeup gas flow (nitrogen) of 8.2 ml min^-1^. Authentic standards (GC grade) were used for qualitative and quantitative analysis.

### Guard cell properties and guard cell starch content

To determine selected GC parameters and starch contents, we used epidermal peels from plants grown under standard conditions in the middle of the photoperiod (6 h illumination). Briefly, nail polish was applied on the abaxial side of the leaves and incubated for 10 min. Further, the epidermis was separated from leaves (6 replicates per line), putted on a slide with water and covered with a coverslip. The prepared slides were subjected to microscopic analysis on a U-LH100HG microscope (Olympus Corporation, Japan) to determine selected GC properties using the manufacturers software. The GC starch content was quantified in epidermal peels (4 biological replicates per genotype, 10 GC per stained peel) as described in Flütsch et al. (2018) following propidium iodide staining. Images were acquired using the Keyence BZ-X800 fluorescence microscope (Keyence Deutschland GmbH, Neu-Isenburg, Germany) equipped with Plan Fluorite 20-100X LD PH objective (100x magnification). Fluorescence was visualized using a BZ-X filter GFP cube (exposure time 1/70 s). Images were captured with the BZ-X800 Viewer software.

### Statistical Analysis

Statistical differences were determined by ANOVA (SPSS Statistics 27, IBM). We used the term significant here if the change in question has been confirmed to be significant at the level of \**p* < 0.05. The Metaboanalyst 6.0 platform (Ewald et al., 2024) was sued to analyze and display the metabolic data.

## Results

### Isolation of guard cell specific Arabidopsis photorespiration mutants

To investigate whether photorespiration is active in guard cells (GC) and involved in stomata movements, we produced transgenic Arabidopsis lines with modified expression of glycine decarboxylase H-protein (GDC-H). GDC-H was chosen based on previous research, demonstrating GDC is key in controlling carbon flux through photorespiration, influencing photosynthesis and growth (Timm et al., 2012a; Simkin et al., 2017; López-Calcagno et al., 2019). To increase (overexpression) and decrease (antisense suppression) *GDC-H* expression in GC, the full-length sequence encoding the *Flaveria pringlei* GDC H-protein (*FpGDC-H*) was cloned into the plant transformation vector pGREEN0229 (http://www.pgreen.ac.uk/). This was done under the control of the GC-specific *GC1* promoter (Yang et al., 2008; Wang et al., 2014) and the CaMV 35S terminator, in both, sense and antisense orientations (Supp. Fig. S1A, B). These constructs were transformed into Arabidopsis wildtype plants. Phosphinotricine (BASTA)-resistant plants were PCR-verified, propagated to stable T3 generations, and used for characterization (Supp. Fig. 1C). Expression of the transgene was confirmed at the mRNA level (Supp. Fig. S1D), followed by tissue-specific analysis of GDC-H protein abundance through immunoblotting. As shown in Fig. 1A and Supp. Fig. 1E, GDC-H remained virtually unchanged in mesophyll cells (MC). However, in GC preparations, we observed increased (∼20-22%) or decreased (∼16-18%) GDC-H abundance in three overexpression (SL2, SL4, and SL7) and three antisense lines (AL4, AL5, and AL9), respectively (Fig. 1B, Supp. Fig. S1E). Notably, GC-specific alterations did not affect the whole-leaf GDC-H abundance (Supp. Fig. S1E), confirming the specificity of the approach.

**Figure 1.**
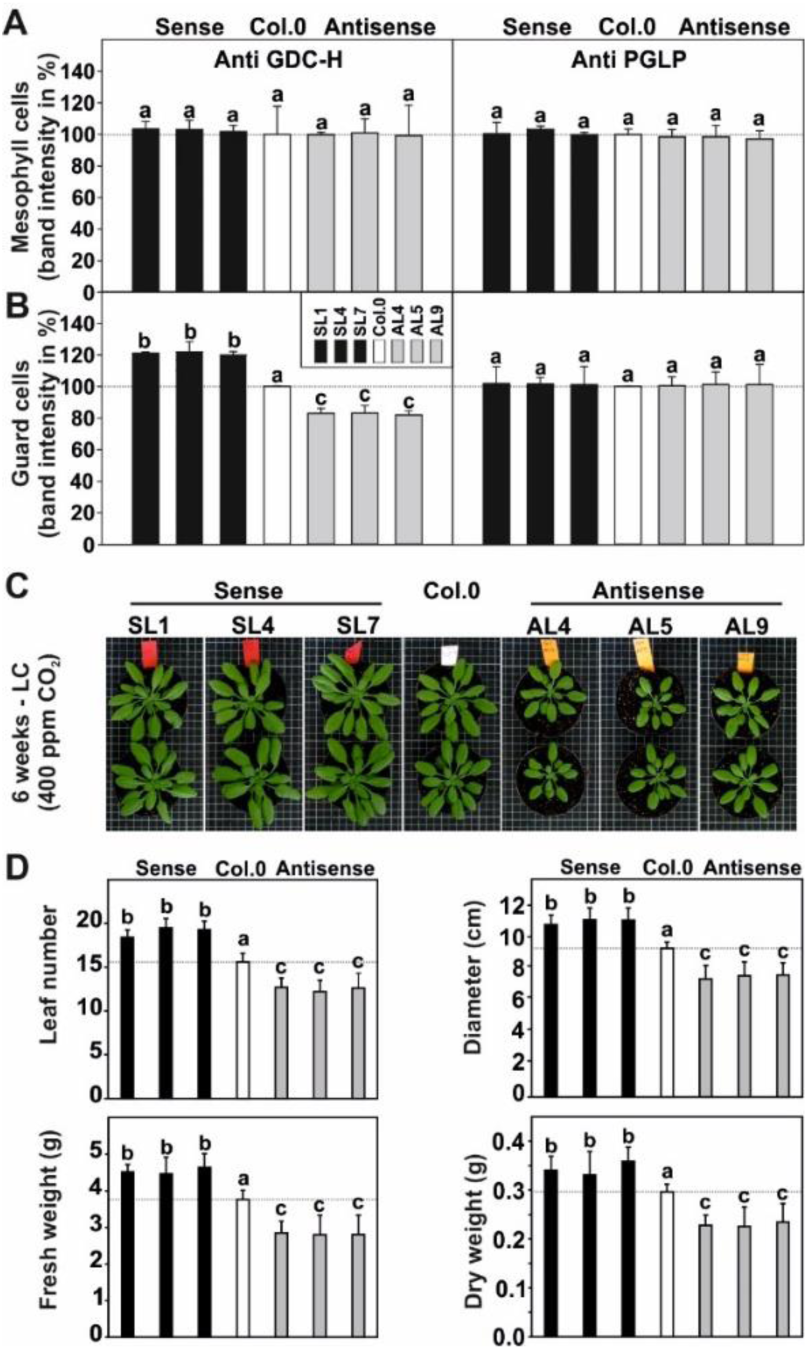
GDC-H protein expression, phenotype and growth of the transgenic, GC-specific *GDC-H* modified lines and the wildtype. Given are GDC-H (left panels) and PGLP1 (right panels) protein abundances in **(A)** mesophyll cell (MC) and **(B)** guard cell (GC) preparations obtained from plants grown in normal air to stage 5.1 (Boyes et al., 2001). Protein expression was quantified by densitometry and expressed in % band intensity (mean ± SD, from 3 individual immunoblots) of signals from the transgenic lines and the wildtype, which was arbitrarily set to 100% (immunoblot-examples are given in Supp. Fig. S1). **(C)** Representative images of plants grown in normal air (LC – low carbon; 400 ppm CO_2_) in a 12/12 h day-/night-cycle (for pictures of plants grown in high CO_2_ and other photoperiods see Supp. Fig. S2). **(D)** Selected growth parameters of all genotypes at the same age. FW/DW ratios: SL1 = 13.3 ± 0.9^a^; SL4 = 13.5 ± 0.8^a^; SL7 = 13.0 ± 0.7^a^; Col.0 = 12.7 ± 0.9^a^; AL4 = 12.5 ± 0.4^a^; AL5 = 12.4 ± 0.5^a^; and AL9 = 11.9 ± 0.8^a^. Given are means ± SD of at least 8 biological replicates. Values that do not share the same letter are significantly different from each other as determined by ANOVA.

### Guard cell specific manipulation of photorespiration correlates with plant growth

Next, we investigated whether modified *GDC-H* expression in GC affects plant growth. To this end, plants were cultivated with sufficient water supply under standard conditions (12/12 h day-night cycle) and analyzed. As shown in Fig. 1C, we observed a correlation between GC-specific GDC-H abundance and growth. Visual comparisons indicated that overexpression of *GDC-H* stimulated, while reduced *GDC-H* expression led to diminished plant growth (Fig. 1C). It is important to note that growth changes were negligible under elevated CO_2_ conditions (3000 ppm), suppressing photorespiration (Supp. Fig. S2A, lower panel). Absolute biomass quantification under standard conditions revealed that total leaf numbers and rosette diameters were significantly increased in overexpression and decreased in antisense lines. These changes in leaf parameters translated to alterations in overall fresh weight (FW) and dry weight (DW), although the FW/DW ratios remained consistent across all genotypes (Fig. 1D). Interestingly, growth effects were consistent across different photoperiods (10/14, 12/12 and 14/10 h day-night cycles, all at 400 ppm CO_2_), however, they appeared somewhat stronger in longer day-night cycles (Supp. Fig. S2B, C).

### Photosynthetic light reactions are unaffected by guard cell modulated photorespiration

Alongside the phenotyping experiments, we characterized the photosynthetic capacity of the transgenic lines and wildtype plants. Further, we used a combination of chlorophyll fluorescence and gas exchange measurements, to distinguish light reaction driven effects from those resulting from altered carbon fixation reactions, eventually occurring in response to changes in stomatal behavior, i.e. stomatal conductance (*g_s_*). No systematic changes emerged during the comparison of a number of PSII and PSI parameters. The maximum quantum yield of PSII (*F_v_*/*F_m_*) and the maximum oxidation of P700 at the PSI reaction center (*P_m_*) were invariable among the studied genotypes under standard growth conditions (Supp. Fig. S3A). These trends were stable over a wide range of light intensities, as the quantum yields of PSII (Y[II]) and PI (Y[I]) did not significantly differ from each other. The same tendencies were observed also when the relative electron transport rates (rETRII and rETRI) were calculated (Supp. Fig. S3B, C). Further, no significant differences were detected in the non-photochemical quenching of PSII (NPQ), the cyclic electron flow around PSI (CET) or the acceptor (Y[NA]) and donor (Y[ND]) side limitation of PSI (Supp. Fig. S3D, E).

### Increased guard cell GDC-H expression improves light-dependent CO_2_ assimilation

Photosynthetic gas exchange parameters provided a different picture. First, we assessed the light acclimation capability of the transgenic lines and the wildtype through measurements of light response curves. As summarized in Fig. 2 (numerical values in Supp. Tab. S1), we observed significantly increased net CO_2_ assimilation (*A_N_*), stomatal conductance (*g_s_*) and transpiration (*E*) in the overexpression plants, whilst all three measurements were lower in the antisense lines compared to the wildtype. Interestingly, changes in *A_N_* were strongly dependent on *g_s_* as revealed by correlation analysis (Fig. 2A-D). The estimation of the maximum photosynthetic rate (*A_max_*) from the light response curves was significantly greater in the overexpressors, whilst decreases were observed in the antisense lines compared to the wildtype. Similarly, initial slopes of the light response curves (*α*_p_) were accelerated in overexpressor, but significantly unchanged in the suppressor lines compared to the wildtype controls (Supp. Tab. S2). Interestingly, higher and lower *g_s_* in the transgenic plants also affect the intracellular CO_2_ concentrations (*C_i_*), which were in- and de-creased in the overexpression and antisense lines, respectively (Fig. 2E). The intrinsic water use efficiency (WUE_int_) did not significantly vary among the studied genotypes (Supp. Tab. S1).

**Figure 2.**
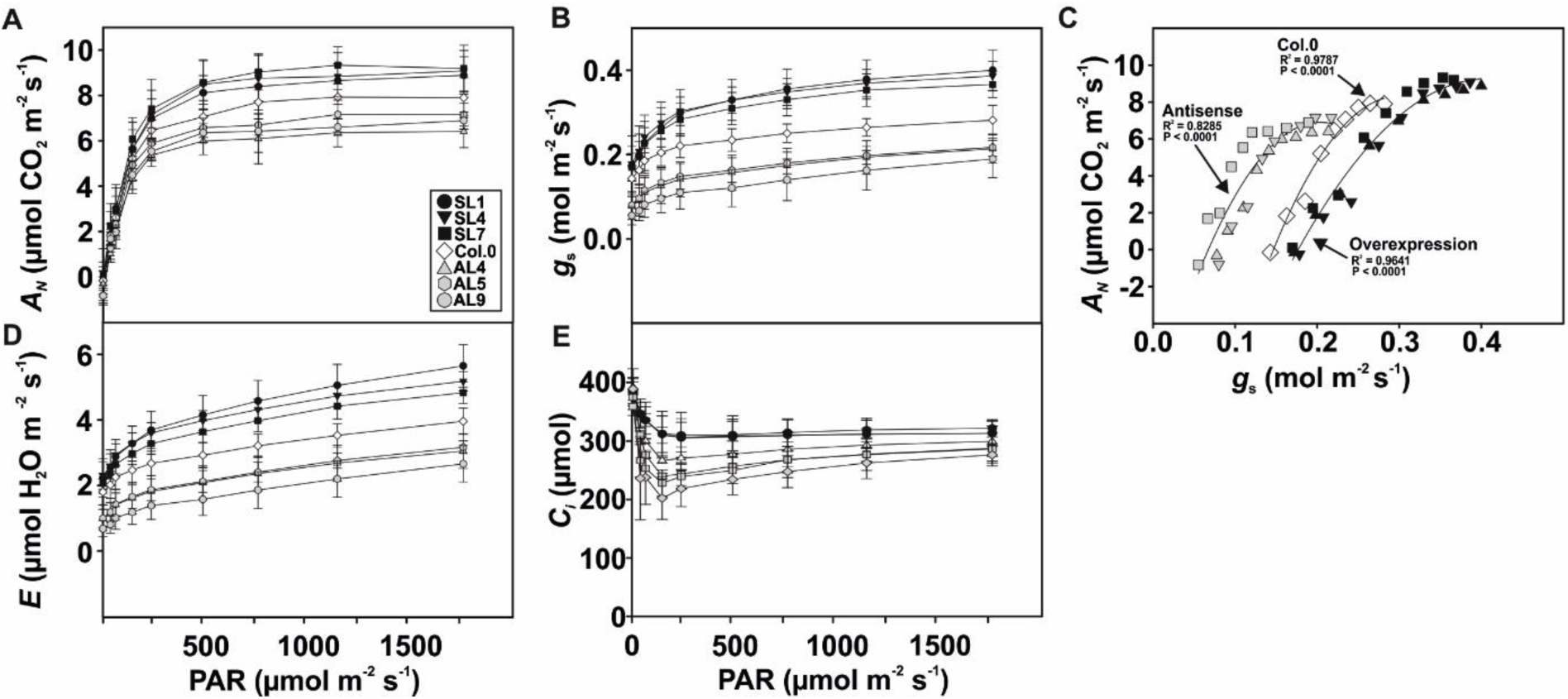
Light response curves of the transgenic, GC-specific *GDC-H* modified lines and the wildtype. Given are selected leaf-gas exchange parameters as functions of varying light intensities determined of plants at stage 5.1 (Boyes et al., 2001) grown under standard conditions. Given are: **(A)** net CO_2_ assimilation rate (*A_N_*); **(B**) stomatal conductance (*g_s_*), **(C)** correlation plot of *A_N_* and *g_s_*, **(D)** transpiration rate (*E*), and, **(E)** intracellular CO_2_ concentration [*C_i_*]. Shown are means ± SD of at least 6 biological replicates. Further parameters, all numerical values and statistical evaluation is provided in Supp. Table S1.

### Guard cell GDC-H expression specifically impacts on O_2_-dependent leaf gas exchange

Given our transgenic approach directly targeted photorespiration, we next measured CO_2_-response curves at three different O_2_ concentrations (3, 21, and 40%) to follow responses with various photorespiratory flux requirements. Net CO_2_ assimilation rates (*A_N_*) and CO_2_ compensation points (*Γ*) were significantly invariant among the genotypes at 3% O_2_, suppressing photorespiration (Fig. 3A, B). However, at 21% O_2_, *A_N_* was elevated (up to ∼16.3%) or lowered (up to ∼18.9%) in the overexpression and antisense lines, respectively (Fig. 3A). In agreement, *Γ* was significantly decreased (up to ∼13.6%) in the overexpression and increased (up to ∼13.3%) in the antisense lines (Fig. 3B). These tendencies were pronounced at 40% O_2_, as the overexpressors displayed increased *A_N_* (up to ∼22%) and a decrease in *Γ* (up to ∼19%), whilst *A_N_* of the antisense lines was reduced (between 26.2-to-36.4%) alongside with a significant rise of *Γ* (up to ∼15.0%) (Fig. 3B). The deduced values of O_2_-inhibition of *A_N_* largely supported the described tendencies, given the overexpression lines were less and antisense lines more inhibited by high O_2_ compared to the wildtype (Fig. 3A). We also calculated *γ*, the slope of the *Γ*-versus-O_2_ concentration function, representing a measure of the photorespiratory CO_2_-release. This parameter was significantly lower in overexpression lines and elevated in the antisense suppressors (Fig. 3B). Related to stomatal functioning, we assessed stomatal conductance (*g_s_*) and transpiration rates (*E*) in the same experiment. Similar to the above trends, *g_s_* and *E* were increased in overexpression and decreased in antisense lines at all O_2_ concentrations (Fig. 3C, D). The only exception was statistically invariant *g_s_* at 3% O_2_ suppressing photorespiration (Fig. 3C).

**Figure 3.**
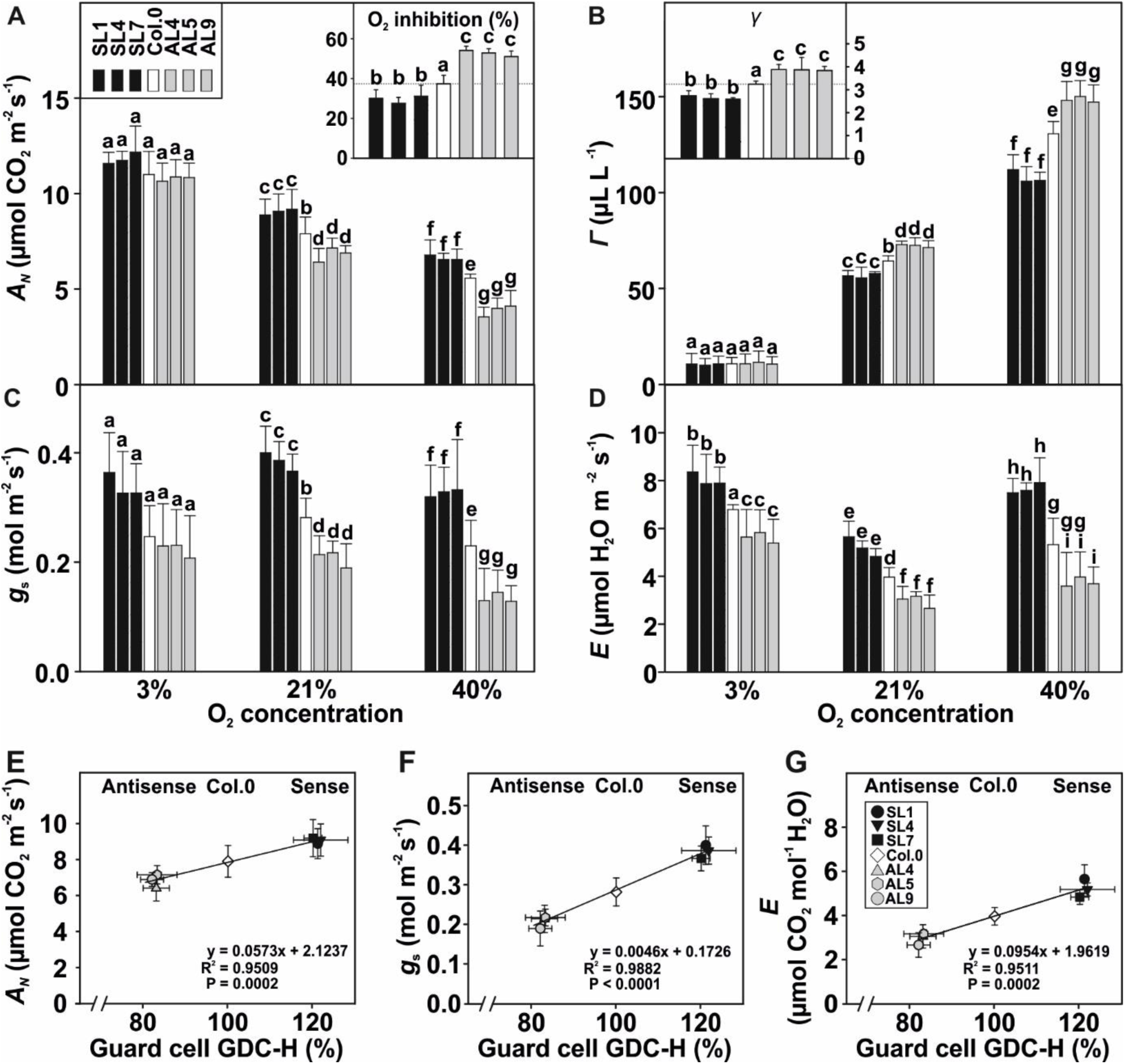
Oxygen-dependent gas exchange parameters of GC specific GDC-H modulated lines and the wildtype. Displayed are selected leaf-gas exchange parameters at three different O_2_ concentrations (3, 21, and 40%, balanced with N_2_) determined using all genotypes grown under standard growth conditions to stage 5.1 (Boyes et al., 2001). Given are: **(A)** net CO_2_ assimilation rates (*A_N_*) and O_2_-inhibition of *A* (insert, overexpression lines were less [-16.6 to -26.2%], whilst antisense lines were more [+36.6 to +34.8%] inhibited at increased O_2_); **(B**) CO_2_ compensation points (*Γ*) and *γ*, slopes of *Γ* – versus - O_2_ curves, (insert); **(C)** stomatal conductance (*g_s_*); and, **(D)** transpiration rates (*E*). Shown are means ± SD of at least 6 biological replicates. Values that do not share the same letter are significantly different from each other as determined by ANOVA. Correlation analysis between GC GDC-H protein expression and **(E)** *A_N_*, **(F)** *g_s_*, and **(G)** *E*, measured in 21% O_2_.

To verify if the measured alterations in leaf-gas exchange parameters correlate with the photorespiratory flux in GC, we plotted their GDC-H amounts against selected photosynthetic parameters. Supporting our hypothesis, we found a strong positive correlation of between GC GDC-H protein expression and *A_N_*, *g_s_* and *E* measured under the ambient air O_2_ concentration of 21%. Hence, all three parameters were lower in the antisense and higher in the overexpression mutants compared to the wildtype (Fig. 3 E-G).

### Optimized GC photorespiration increases accumulation of leaf-carbohydrates

To determine if improved *g_s_* and photosynthesis facilitates accumulation of photosynthates, eventually stimulated growth, we quantified total leaf-abundances of selected carbohydrates. Three soluble sugars (sucrose, glucose, and fructose) and transitory starch were measured in standard condition grown plants. As summarized in Table 1, soluble sugars were significantly elevated in the overexpression, but lower in the antisense lines compared to the wildtype. Further, transitory starch measured in the same material followed the described pattern, as it was increased in the overexpressors and reduced in the antisense lines (Table 1).

**Table 1.** Soluble sugars and starch in leaves of the transgenic lines and the wildtype. Plants were grown under environmental controlled standard conditions in normal air (400 ppm CO_2_) to growth stage 5.1 (Boyes et al., 2001). Leaf material was harvested at end of the day (EoD – 11 h illumination) and used for quantification of selected soluble sugars by gas chromatography. Starch was measured spectrophotometrically essentially using the same material. Values are means ± SD (n = 6) and are given in µg per g DW^-1^. Values that do not share the same letter are significantly different from each other as determined by ANOVA.

| Compound |  |  |  |  |
| --- | --- | --- | --- | --- |
| Genotype | Sucrose | Fructose | Glucose | Starch |
| SL1 | **5.88 ± 0.89<sup>a</sup>** | **2.88 ± 0.53<sup>b</sup>** | **17.76 ± 3.16<sup>b</sup>** | **152.32 ± 12.81<sup>b</sup>** |
| SL4 | **5.36 ± 0.52<sup>a</sup>** | **2.73 ± 0.18<sup>b</sup>** | **16.15 ± 1.19<sup>b</sup>** | **159.53 ± 29.32<sup>b</sup>** |
| SL7 | **5.18 ± 0.29<sup>a</sup>** | **2.68 ± 0.57<sup>b</sup>** | **14.12 ± 1.53<sup>b</sup>** | **145.62 ± 10.85<sup>b</sup>** |
| Col.0 | 4.60 ± 0.28<sup>b</sup> | 1.90 ± 0.26<sup>a</sup> | 8.52 ± 2.35<sup>a</sup> | 117.54 ± 18.48<sup>a</sup> |
| AL4 | **3.89 ± 0.45<sup>c</sup>** | **1.25 ± 0.24<sup>c</sup>** | **4.37 ± 0.78<sup>c</sup>** | **73.37 ± 18.63<sup>c</sup>** |
| AL5 | **3.95 ± 0.34<sup>c</sup>** | **1.18 ± 0.20<sup>c</sup>** | **3.48 ± 1.55<sup>c</sup>** | **77.26 ± 15.16<sup>c</sup>** |
| AL5 | **3.60 ± 0.41<sup>c</sup>** | **1.24 ± 0.21<sup>c</sup>** | **3.99 ± 2.58<sup>c</sup>** | **72.67 ± 12.42<sup>c</sup>** |

### Primary metabolism in the GC specific transgenic lines responds with specific shifts associated with 3PGA and amino and organic acid metabolism

Simultaneous to carbohydrate measurements, we quantified 35 further primary metabolites, including four photorespiratory intermediates (2-PG, glycine, serine and 3-PGA), through liquid chromatography coupled to tandem mass spectrometry (LC-MS/MS). Only a few consistent, mutant specific, alterations were seen in leaves at end of the day from plants grown under standard conditions (see Supp. Table S3). However, overexpression lines generally tend to accumulate soluble amino acids (significant in SL1 and SL7), whilst organic acids were decreased. By contrast, antisense lines displayed somewhat opposite tendencies, with soluble amino acids being decreased (significant in AL5 and AL9) and organic acids were present in similar amounts as in the wildtype (Fig. 4A). Related to photorespiration, we found glycine and serine amounts statistically invariant among the genotypes and only slight decreases in 2-PG in the overexpressors. Interestingly, 3-PGA followed the same pattern described for carbohydrates (Table 1), as its contents were increased in overexpressors and decreased in antisense lines (except for AL9) (Fig. 4B).

**Figure 4.**
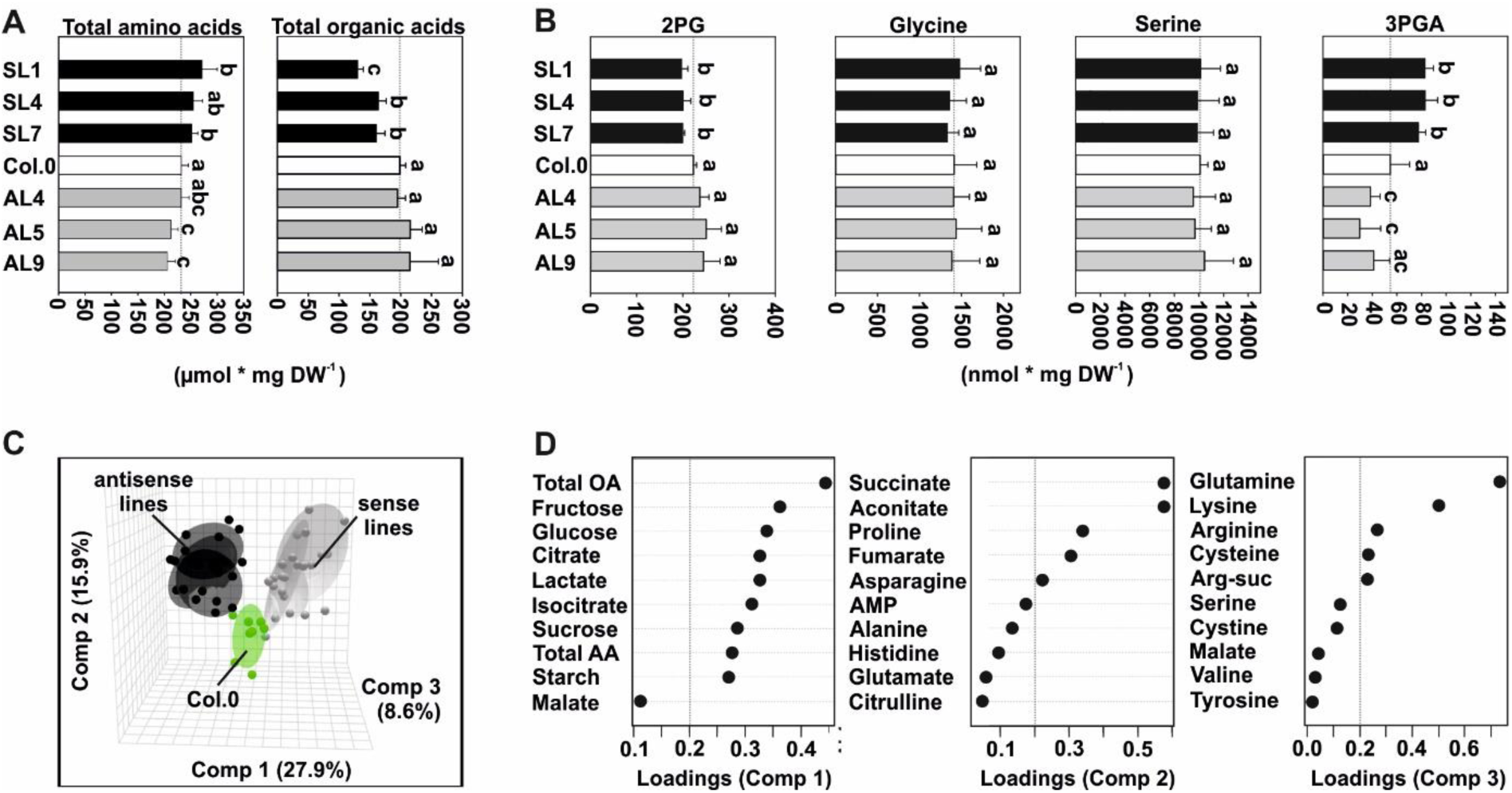
Overview of metabolite responses in GC specific GDC-H modulated lines and the wildtype. Plants were grown under environmental controlled standard conditions in normal air (400 ppm CO_2_) to growth stage 5.1 (Boyes et al., 2001). Leaf-material was harvested at end of the day (EoD – 11 h illumination) and used for quantification of primary metabolites by LC-MS/MS analysis. Given are **(A)** total soluble amino acid and organic acid contents, **(B)** selected intermediates of photorespiration, **(C)** principal component analysis (PCA) of all metabolite data and **(D)** loadings of the first three components driving the cluster separation during PCA. Loadings exceeding the cutoff (± 0.2) are presented bold. Metabolite contents are means ± SD (n > 6) and are given in µmol (A) or nmol (B) per g DW^-1^, respectively. Values that do not share the same letter are significantly different from each other as determined by ANOVA. Abbreviations: Comp – component, total OA – total organic acids, total AA – total amino acids, AMP – adenosine monophosphate, and Arg-suc – L-argininosuccinic acid.

To gain a general, comparative overview on the metabolic changes, we analyzed the full data set (GC, starch and LC-MS/MS data) by principal component analysis (PCA). As shown in Fig. 4C, metabolites in the different genotypes formed three different clusters, namely, the wildtype (1, green), the antisense (2, dark grey) and the overexpression (3, light grey). Formation of the clusters was mainly driven in the plain of PC1 (27.9%) and PC2 (15.9%), whilst PC3 (8.6%) was only responsible for a small separation of the data (Fig. 4C). A closer view on the factors driving the differentiation revealed high positive loadings of the total organic and amino acid counts, carbohydrates (glucose, fructose, sucrose and starch) and the individual organic acids citrate, lactate and isocitrate for PC1. Separation along PC2 was mainly driven by organic (succinate, aconitate and fumarate) and amino acids (proline and asparagine). Finally, PC3 separation was due to the amino acids glutamine, lysine and arginine (Fig. 4D; Supp. Tab. S4).

### Stomata count and size are unaffected, whilst GC starch showed a positive correlation with GDC-H protein expression

To discover if modified GC photorespiration caused alterations in stomata properties (e.g. number and size), we analyzed epidermal peels by microscopy. As shown in Table 2, no significant changes were observed in stomatal density and index. Further, stomata were invariant among all genotypes with regards to either, length, width or area (Table 2).

**Table 2.** Overview on stomatal parameters of GC specific GDC-H modulated lines and the wildtype. For microscopic determination of stomatal parameters, we used plants grown under standard conditions to growth stage 5.1 (Boyes et al., 2001). Given are stomatal density, stomata length, stomata width, stomatal area and stomatal index, respectively. Shown are means ± SD of at least 4 biological replicates, with measurements of at least 10 stomata per leaf (40 stomata in total). Values that do not share the same letter are significantly different from each other as determined by ANOVA.

| Parameter |  |  |  |  |  |
| --- | --- | --- | --- | --- | --- |
| Genotype | Stomatal density<br>(mm<sup>-2</sup>) | Long axis<br>(μm) | Short axis<br>(μm) | Stomatal area<br>(μm<sup>-2</sup>) | Stomatal index<br>(mm<sup>2</sup> mm<sup>-2</sup>) |
| SL1 | 142.48 ± 8.85<sup>a</sup> | 20.99 ± 1.40<sup>a</sup> | 14.66 ± 1.48<sup>a</sup> | 966.22 ± 144.89<sup>a</sup> | 0.14 ± 0.01<sup>a</sup> |
| SL4 | 157.76 ± 19.56<sup>a</sup> | 21.13 ± 1.51<sup>a</sup> | 14.68 ± 0.61<sup>a</sup> | 975.77 ± 90.72<sup>a</sup> | 0.15 ± 0.02<sup>a</sup> |
| SL7 | 167.14 ± 32.48<sup>a</sup> | 20.04 ± 1.47<sup>a</sup> | 13.72 ± 0.78<sup>a</sup> | 864.41 ± 79.84<sup>a</sup> | 0.14 ± 0.03<sup>a</sup> |
| Col.0 | 148.10 ± 20.30<sup>a</sup> | 20.34 ± 0.63<sup>a</sup> | 14.33 ± 1.53<sup>a</sup> | 917.31 ± 106.78<sup>a</sup> | 0.13 ± 0.01<sup>a</sup> |
| AL4 | 156.03 ± 27.80<sup>a</sup> | 20.22 ± 1.88<sup>a</sup> | 13.82 ± 1.43<sup>a</sup> | 882.94 ± 143.41<sup>a</sup> | 0.14 ± 0.02<sup>a</sup> |
| AL5 | 168.72 ± 37.97<sup>a</sup> | 20.65 ± 1.24<sup>a</sup> | 14.08 ± 0.77<sup>a</sup> | 913.27 ± 67.03<sup>a</sup> | 0.15 ± 0.04<sup>a</sup> |
| AL9 | 155.31 ± 12.84<sup>a</sup> | 20.30 ± 0.93<sup>a</sup> | 14.25 ± 0.93<sup>a</sup> | 912.00 ± 124.10<sup>a</sup> | 0.14 ± 0.01<sup>a</sup> |

Given previous research revealed that GC starch amounts are key for their metabolism, i.e. rapid movements of stomata (Flütsch et al., 2020; Dang et al., 2024), we also analyzed the starch content of GC (Fig. 5A) in the same plant material. Interestingly, we observed a positive correlation between the GDC-H protein expression and starch amounts in GC. More precisely, all overexpression lines were significantly increased (up to 26.1%), whilst the antisense suppressors were decreased (up to -23.4%) in GC starch at mid of the day (Fig. 5B-C). Collectively, the data suggest that stomatal aperture and their starch content, but not their size and amount, responded to manipulation of GC photorespiration.

**Figure 5.**
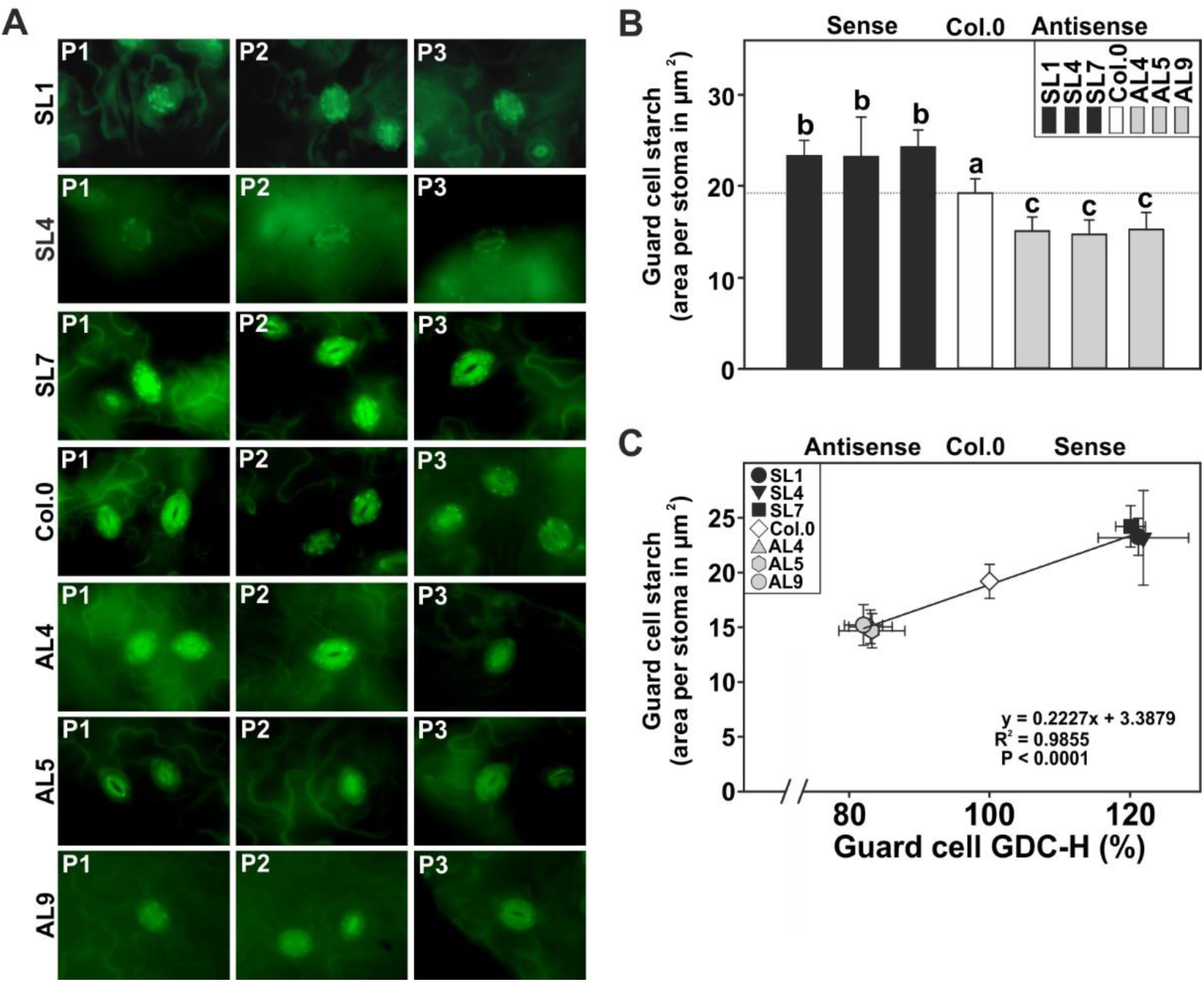
GC starch quantification in GC specific GDC-H mutant lines and the wildtype. For microscopic determination of GC starch accumulation patterns, we used plants grown under standard conditions to growth stage 5.1 (Boyes et al., 2001). Given are **(A)** three representative microscopic images of each genotype following starch staining, **(B)** GC starch accumulation, and, **(C)** correlation analysis between GC GDC-H protein expression and GC starch contents. Shown are means ± SD of at least 4 biological replicates, with at least 10 stomata per leaf [Σ > 40]. Values that do not share the same letter are significantly different from each other as determined by ANOVA.

## Discussion

The important role of photorespiration in enabling efficient photosynthesis at atmospheric O_2_ levels is generally accepted (Bauwe et al., 2012; Busch, 2020). However, specific implications for stomatal behavior, especially GC metabolism, remained unexplored due to the absence of GC-specific mutants (Lemonnier and Lawson, 2024). Earlier work showed that Arabidopsis wildtype plants shifted from high-to-low CO_2_, to induce photorespiratory activity, responded with only a few transcriptional changes, interesting these were mainly genes related to drought stress. This is likely a consequence of reduced CO_2_ availability, prompting enhanced stomatal opening and potentially increased water loss (Eisenhut et al., 2017). Interestingly, when mesophyll cell (MC) photorespiration is genetically impaired, this response pattern was abolished. Such mutants display reduced stomatal conductance (*g_s_*), transpiration rates and stronger transcriptional reprogramming than in wildtype plants. These changes included, among others, a marked reduction in phototropin (*PHOT1* and *PHOT2*) expression, which are essential for light-dependent stomatal opening (Eisenhut et al., 2017) and in particular the species specific stomatal blue light response (Vialet-Chabrand et al.,2021). Collectively, these findings pointed us to the hypothesis that active photorespiration is required for stomatal movements in response to changes in external CO_2_. As earlier work provided evidence for CO_2_ fixation and photorespiration in GC (Reckmann et al., 1990; Cardon and Berry, 1992; Lemonnier and Lawson, 2024), there is a need for further research to unravel potential interactions between photosynthesis, photorespiration and stomatal behavior. Moreover, we speculated that MC photorespiration may directly affect GC metabolism, and, possibly *vice versa*, via contributing to the *C_i_*-dependent regulatory mechanism coordinating *g_s_* and photosynthesis.

Through GC-specific genetic manipulation of the key photorespiratory enzyme glycine decarboxylase (GDC) H-protein (Timm et al., 2012a), we demonstrated that photorespiration is active in GC and serves as a yet unidentified component contributing to optimal stomatal movements and metabolism in Arabidopsis leaves in current atmospheric air. This conclusion aligns with our observation that GC-specific manipulation of *GDC-H* expression impacts on biomass accumulation (Fig. 1, Supp. Fig. 2). The specificity of this manipulation, i.e. altered expression only in GC (Fig. 1, Supp. Fig. 1), supports the previously raised hypothesis that photorespiration is present and fully active in GC, and demonstrates the significant role in supporting stomata functions. It is noteworthy stating biomass changes were specifically attributable to altered photorespiratory fluxes in GC which, in turn, impact on stomatal behavior. This agrees with the fact that transgenic plants were visually indistinguishable under high CO_2_ conditions, suppressing photorespiration (Supp. Fig. 2A), and therefore minimizing any significant impact of photorespiratory manipulations. The statement is consistent with observations resulting from studies on classic photorespiratory mutants, which are lethal in normal air but fully recoverable in high CO_2_ on the phenotypical and physiological level (Sommerville, 2001; Timm et al., 2012b; Eisenhut et al., 2017). Further, biomass alterations in GC (this study) and whole-leaf (e.g. Queval et al., 2007) photorespiratory mutants were pronounced under longer days (Supp. Fig. S2), which can be explained by the longer necessity for increased photorespiratory capacity with extended illumination, during which more toxic pathway intermediates potentially accumulate. Therefore, genetic modifications to photorespiratory pathway exert a greater impact on plants grown under long-day conditions.

Our photosynthetic characterization revealed that GC-specific manipulation of photorespiration did not significantly affect MC photosynthetic light reactions (Supp. Fig. S3), as expected. However, GC-specific overexpression of *GDC-H* stimulated net CO_2_ assimilation rates (*A_N_*) across a wide range of light intensities (Fig. 2A) driven by increasing *g_s_* and the removal of diffusional constraints on CO_2_ uptake indicted by the greater intracellular CO_2_ concentrations (*C_i_*) (Fig. 2B-E). Notably, such physiological responses occurred in the absence of alterations in stomata amount and size (Table 2). This is in line with other studies that have manipulated GC, positively affecting *g_s_* and *A_N_*, with unchanged stomata characteristics (Wang et al., 2014). Notwithstanding, our findings suggest a direct correlation between GC photorespiratory flux and *g_s_*. Mechanistically, this response could result from changes in photorespiratory flux signaling varying CO_2_ availability within GC, which induces changes in aperture to supply more or less CO_2_ for mesophyll photosynthesis. Enhanced mesophyll photosynthesis due to greater *g_s_* in the *GDC-H* over-expression lines supports greater carbohydrate biosynthesis, stimulating growth. Both improved photosynthesis and increased accumulation of photosynthates (e.g., 3-PGA, sucrose, and transitory starch), alongside enhanced growth, were observed in these lines. Conversely, antisense suppression of *GDC-H* led to the opposite effects (Fig. 1–4, Table 1). Similar outcomes have been reported for leaf-specific manipulations of photorespiratory genes, including *GDC* (H- and L-protein) and *PGLP1* (Timm et al., 2012a; 2015; Flügel et al., 2017; Simkin et al., 2017; López-Calcagno et al., 2019). These effects were rationally explained by alleviating negative feedback inhibition of photorespiratory intermediates on the CB-cycle, thereby improving RuBP regeneration and enhancing carbon assimilation and export (reviewed in Bauwe, 2018; Timm and Hagemann, 2020). However, since gene expression driven by promoters like *ST*-*LSI* or *35S* is not fully restricted to MC, it remains uncertain whether the observed effects in these studies result exclusively from MC-specific manipulation of photorespiration or whether GC photorespiratory metabolism and behavior have also contributed toward those changes. Our findings that GC-specific photorespiration plays a significant role in regulation of stomatal movements could be due to the lowering of metabolic inhibitions on the CB-cycle directly in GC and improved CO_2_-fixation in these cells that provides osmotic for increase aperture. The improved photosynthesis in the overexpressors could potentially signal a higher GC CO_2_ demand (i.e. lowered *C_i_*), prompting stomata to open, with the opposite occurs in the antisense lines, characterized by lower photorespiratory flux and impaired CB-cycle activity. Alternatively, the changes in metabolic fluxes in both photorespiration and photosynthesis in the guard cells could also influence GC electron transport rates (Lawson et al., 2002; 2003), which in turn would alter the redox state of the PQ pool, which has been suggested to play a role in regulating *g_s_* in line with mesophyll photosynthetic demands (Lawson et al., 2008; Busch, 2014; Kromdijk et al., 2020; Taylor et al., 2024). In addition to examining light acclimation of photosynthesis, photosynthetic parameters in response to changes in external O_2_ concentrations were profiled, primarily to manipulate leaf photorespiration on a short-term basis. Notably, our measurements revealed that GC-specific manipulation of photorespiration significantly influenced the O_2_ susceptibility of the transgenic plants (Fig. 3). Specifically, increasing the photorespiratory flux in GC reduced the inhibitory effects of high O_2_ on photosynthesis, likely by mitigating the impact of increased Rubisco oxygenation due to improved CO_2_ availability via greater *g_s_* (Fig. 3C, F). Conversely, reducing the photorespiratory flux resulted in impaired photosynthetic parameters under the same CO_2_/O_2_ ratios. It is worth highlighting that differences in photosynthetic parameters were negligible under low O_2_ conditions. This finding is consistent with the normalization of growth observed under elevated CO_2_, conditions that drastically diminish the need for efficient photorespiratory metabolism (Fig. 3). Furthermore, our findings agree with recent studies that highlight the utility of altered O_2_ levels as a proxy for modulating photorespiratory flux, even in the absence of genetic perturbations to the pathway (Fu et al., 2023; Smith et al., 2023).

Alongside with specific changes in leaf-carbohydrate accumulation patters (Table 1), we observed only a few metabolic shifts in the transgenic lines through LC-MS/MS analysis. These measurements revealed that plants mainly responded with a general increase in soluble amino acids and a decrease in organic acids following GC specific upregulation of the photorespiratory flux, with the opposite holds true for antisense suppressors (Fig. 4). Hence, it seems there is a correlation with the elevated photosynthesis and the increased availability of carbon building blocks that is potentially used to support higher nitrogen assimilation rates to support amino acid biosynthesis to facilitate protein biosynthesis and increased plant growth (Fig. 1, Supp. Fig. S2). This hypothesis is in line with experimental support from other research that pioneered a correlation between the photorespiratory flux, adjusted by external CO_2_ availability, and nitrogen assimilation (Rachmilevitch et al., 2004; Bloom et al., 2010). Alternatively, increased amino acid-to-organic acid ratio can be explained by higher ammonia release in response to GDC activity upregulation.

Finally, and more importantly, simultaneous to the higher whole leaf-carbohydrate status in the overexpressors (Table 1) we determined increased starch accumulation in GC of the overexpressors, whilst lower amounts were present in the antisense mutants (Fig. 5). This observation could also explain altered stomatal behavior in the transgenic lines as GC starch amount were reported to be of high importance for stomatal movements (Flütsch et al., 2020; Dang et al., 2024). Based on their findings, the authors suggest a higher availability in GC starch results in an improved glucose availability, supporting rapid stomatal movements (Flütsch et al., 2020). Nevertheless, it remains an open question if starch synthesis in GC is directly affected in response to GC specific photorespiratory flux manipulations or if this is a result of the increased MC photosynthesis and carbohydrate biosynthesis. The latter argument is in favor with the general assumption that GC starch is mainly synthesized using carbon building blocks imported from the MC and further agrees with research on Arabidopsis sucrose synthases (Daloso et al., 2016; Piro et al., 2023).

The findings of this study are summarized in the tentative model illustrated in Fig. 6. This model proposes that changes in external CO_2_ availability are detected and translated by the guard cell photorespiratory flux in response to alterations in *C_i_*, which modulates photosynthetic activities, electron transport or an other unknown metabolic process within these specialized cells. Consequently, stomatal movements are adjusted according to the availability of energy resources, including the amounts of GC starch (Fig. 5) and or sense changes in CO_2_ to which they respond. Mechanistically, variations in photorespiratory flux influence CB-cycle activities by alleviating or intensifying negative metabolic feedback on the regenerative branch of the pathway and altering the extent of carbon export from the cycle. This hypothesis is strongly supported by experimental evidence from studies manipulating photorespiratory flux in MC (Timm et al., 2012a, 2015; Flügel et al., 2017). Furthermore, we propose that changes in GC photorespiratory flux, particularly glycine decarboxylation by GDC, directly impact on guard cell *C_i_*, thereby influencing photosynthetic activities. On one hand, the surplus CO_2_ released from mitochondria becomes available for the CB-cycle by reducing the oxygenation reaction of Rubisco (Fig. 6). On the other hand, the increase in mitochondrially driven CO_2_ could, at least partially, be converted to bicarbonate via carbonic anhydrase, facilitating phospho*enol*pyruvate carboxylase (PEPC)-mediated CO_2_ fixation in GC. This process would result in increased energy resources to drive stomatal movements. However, further work is needed to fully elucidate the exact mechanism by which the photorespiratory process specifically in GC influence stomatal behavior.

**Figure 6.**
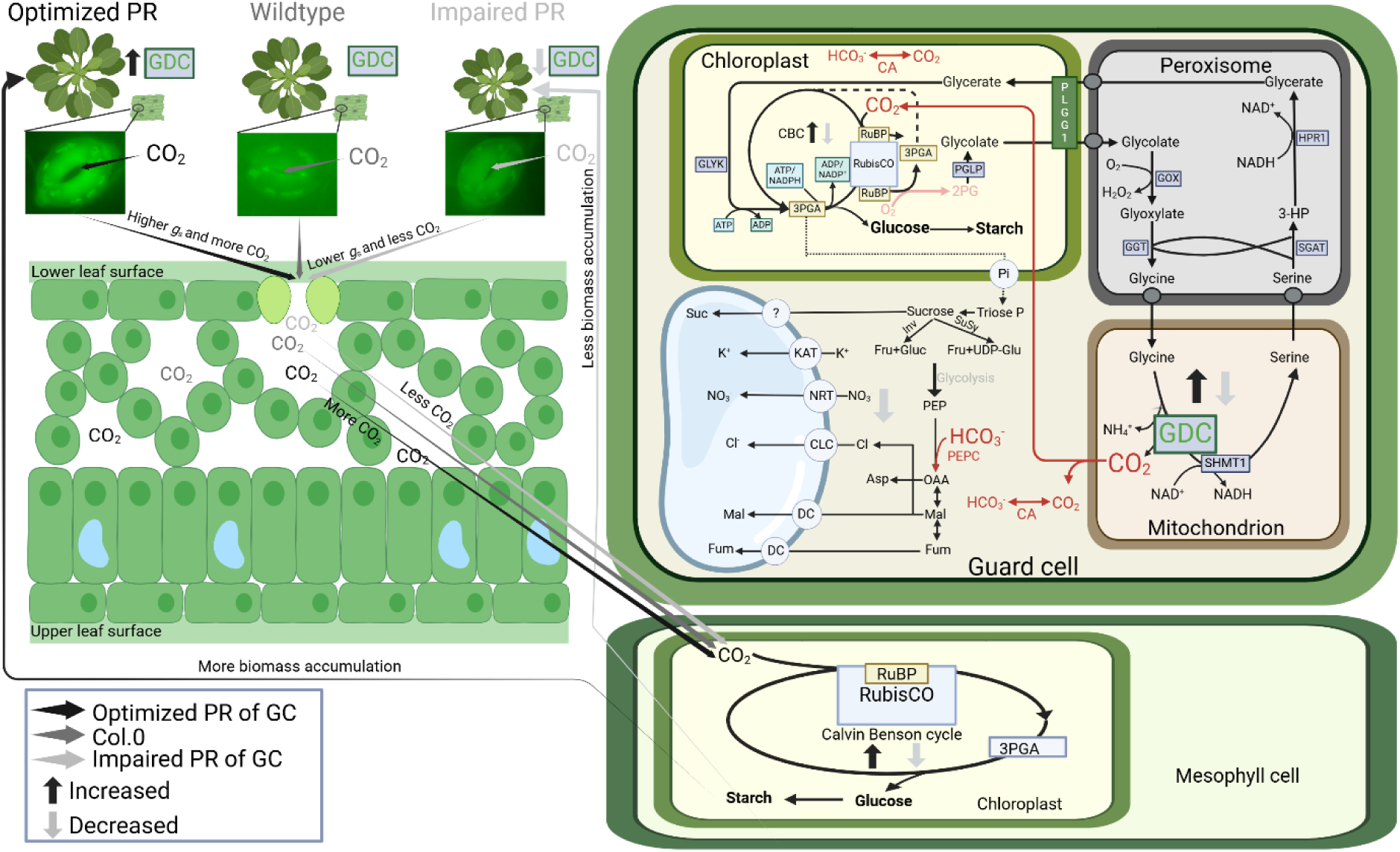
Scheme of current hypothesis on the impact of altered GC photorespiration. Stomata physiologically respond to changes in external CO_2_ availability, in turn affecting internal CO_2_ (*C_i_*). High CO_2_ leads to stomatal closure, whilst low CO_2_ leads to stomatal opening. Changes in *C_i_*, and the resulting movements, can be mimicked through our transgenic intervention, altering the guard cell photorespiratory flux. At one hand, low external CO_2_ enhances the photorespiratory flux (more Rubisco oxygenation), which is similar in the GDC-H overexpression plants, and prompts stomata to be more open (low *C_i_*). On the other hand, high external CO_2_ suppresses the photorespiratory flux (less Rubisco oxygenation), which is comparable with our guard cell GDC-H antisense lines, causing stomata to be more closed (high *C_i_*). Mechanistically, improved or lower (impaired) GC photorespiratory flux leads to alleviated or enhanced negative metabolic feedback onto the CB-cycle, improving or hindering RuBP regeneration. Improved CB-cycle performance lowers *C_i_* and makes GC a stronger CO_2_ sink, enhances stomatal opening (higher *g_s_*), to ultimately provide more CO_2_ for mesophyll photosynthesis. Higher photosynthesis stimulated carbohydrate biosynthesis and plant growth. Impaired or repressed CB-cycle performance lowers CO_2_ requirements, leads to less open stomata (lower *g_s_*) and decreases mesophyll CO_2_. Reduced mesophyll photosynthesis provides less carbon for plant growth. Further, we hypothesize that manipulation of GC GDC activity (glycine decarboxylation) through increased and suppressed GDC-H protein expression has a direct impact on the GC *C_i_*, too. More GDC activity (in overexpression lines, increased glycine decarboxylation) leads to increased, whilst lower GDC activity (in antisense lines, reduced glycine decarboxylation) leads to reduced GC *C_i_*. Changes in GC *C_i_* supports (or reduces) photosynthesis for increased (or decreased) GC starch production, providing more (or less) energy supply for stomatal movements. Changes in CO_2_ affects both, Rubisco and PEPc mediated GC photosynthesis, respectively. The figure was created with bioRender (https://www.biorender.com/). Abbreviations: enzymes; CA - carbonic anhydrase; Inv - invertase; PEPc - phospho*enol*pyruvate carboxylase; PEPk - phospho*enol*pyruvate carboxykinase; RubisCO - ribulose-1,5-biphosphate carboxylase/oxygenase; SuSy - sucrose synthase. Metabolites; 3PGA - 3 phosphoglycerate; Asp - aspartate; Cl^−^-chloride; Fru - fructose; Fum - fumarate; Glu - glutamate; Gluc - glucose; Isoc - isocitrate; K^+^-potassium; Mal - malate; NO ^−^ - nitrate; OAA - oxaloacetate; PEP, phospho*enol*pyruvate; Suc - sucrose; UDP-glu - uridine diphosphate glucose. Transporters; CLC - chloride channel; DC - dicarboxylate carrier; NRT - nitrate transporter; OC - putative oxaloacetate carrier.

## Conclusion and outlook

Based on the findings on the GC-specific manipulation of photorespiration, we propose a novel function of photorespiration for the C3 plant Arabidopsis. It appears that the GC photorespiration is active and crucial for optimal stomatal behavior, including stomatal conductance, and could represent a prerequisite to adapt to variations in the external CO_2_/O_2_ ratios that affect *C_i_* (Fig. 6). It seems reasonably to hypothesize that changes in GC photorespiratory flux could eventually signal the GC CO_2_ demands in response to external CO_2_ availability, in turn communicating with the mesophyll to adjust mesophyll photosynthesis and carbohydrate biosynthesis. However, how alterations in the GC photorespiratory flux are sensed and incorporated into the CO_2_ sensing and signaling cascade remains an open question. In order to resolve this regulation circuit in more detail, future studies of MC and GC specific photorespiratory manipulations in GC CO_2_ signaling mutant backgrounds are required.

## Supporting information

Supplemental data Photorespiration in guard cells

## Acknowledgements

H.S. gratefully acknowledges the scholarship granted by the China Scholarship Council (CSC). We wish to thank Prof. Hendrik Schubert for access to the Dual PAM-100 and Junior Prof’s. Klaus Herburger and Andreas Richter for access to the confocal laser scanning microscope and assistance with the gas chromatography measurements (all at Rostock University). The technical assistance received from Klaudia Michl and Kathrin Jahnke (Rostock University) is highly appreciated. This work was supported by the University of Rostock to M.H. and S.T.

## Author Contributions

S.T. conceived and supervised the project. H.S., and S.T. designed the research. N.S. performed cloning procedures and established the transgenic lines. H.S. performed the research. H.S., T.L., M.H., and S.T. analyzed the data. M.H. provided experimental equipment and tools. S.T. wrote the article, with additions and revisions from H.S., T.L., and M.H. All authors have read and approved the final version of the manuscript.

## Data availability

The data supporting the findings of this study are presented as Figures 1–5 and Table 1 included in the main text and Supplemental Data (Figures S1–S3; Tables S1–S4 associated with this article. Plasmids and transgenic plants generated in this study will be made available upon request to the corresponding author.

## Supplemental data

**Supplemental Figure S1.** Generation and verification of Arabidopsis GC specific *GDC-H* overexpression and antisense lines.

**Supplemental Figure S2.** Phenotype of Arabidopsis GC specific GDC-H modulated lines and the wildtype under different growth conditions.

**Supplemental Figure S3.** Chlorophyll fluorescence parameters of Arabidopsis GC specific GDC-H modulated lines and the wildtype.

**Supplemental Table S1.** Light response curves of the transgenic lines and the wildtype under standard conditions.

**Supplemental Table S2.** Calculated parameters from light response curves numerically given in Suppplemental Table S1.

**Supplemental Table S3.** Abundances of selected intermediates associated with primary metabolism in the transgenic lines and the wildtype under standard conditions.

**Supplemental Table S4.** Loadings of metabolites on the first three principal components (PCs) in leaves of the GC specific GDC-H lines and the wildtype.

**Supplemental Table S5.** Primers used for PCR amplification of genomic DNA and cDNA.

