## Supplemental data Photorespiration in guard cells for "Guard cell photorespiration has a major impact on photosynthesis, growth and stomatal behavior in Arabidopsis"

#### Supplemental Figures

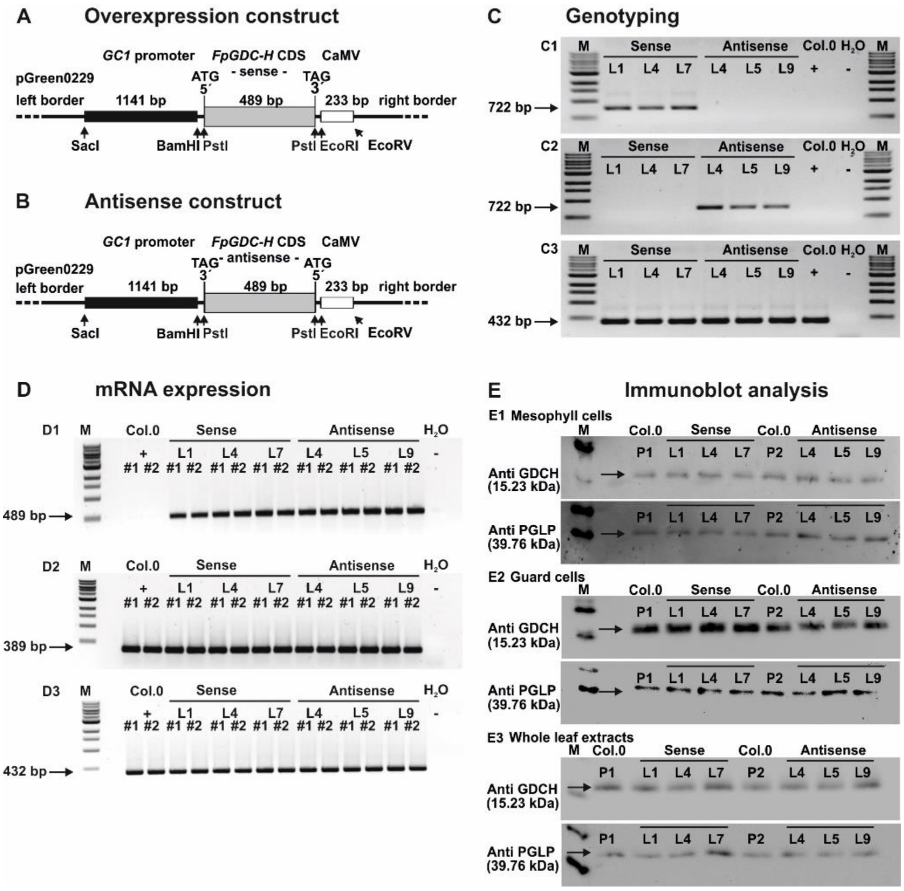


**Supplemental Figure S1.** Generation and verification of Arabidopsis GC specific *GDC-H* overexpression and antisense lines.

Schematic overview of the GC specific *FpGDC-H* **(A)** overexpression and **(B)** antisense constructs. **(C)** PCR verification of the transformed constructs into the genome of transgenic **(C1)** overexpression and **(C2)** antisense lines and the corresponding loading control **(C3)**. **(D)** RT-PCR verification of the full-length *FpGDCH* **(D1)** and the *AtGDCH1* **(D2)** transcripts, in comparison with signals of the constitutively expressed 40S ribosomal protein *S16* gene as loading control **(D3)**. **(E)** Immunoblots of protein extracts from **(E1)** MC, **(E2)** GC and whole leaves **(E3)** using a specific antibody against mitochondrial GDC-H (Timm et al., 2013) and chloroplastidal 2-phosphoglycolate phosphatase 1 (PGLP1; Flügel et al., 2017) as loading control. Note, loading control immunoblots were essentially developed from the same membrane as for GDC-H.

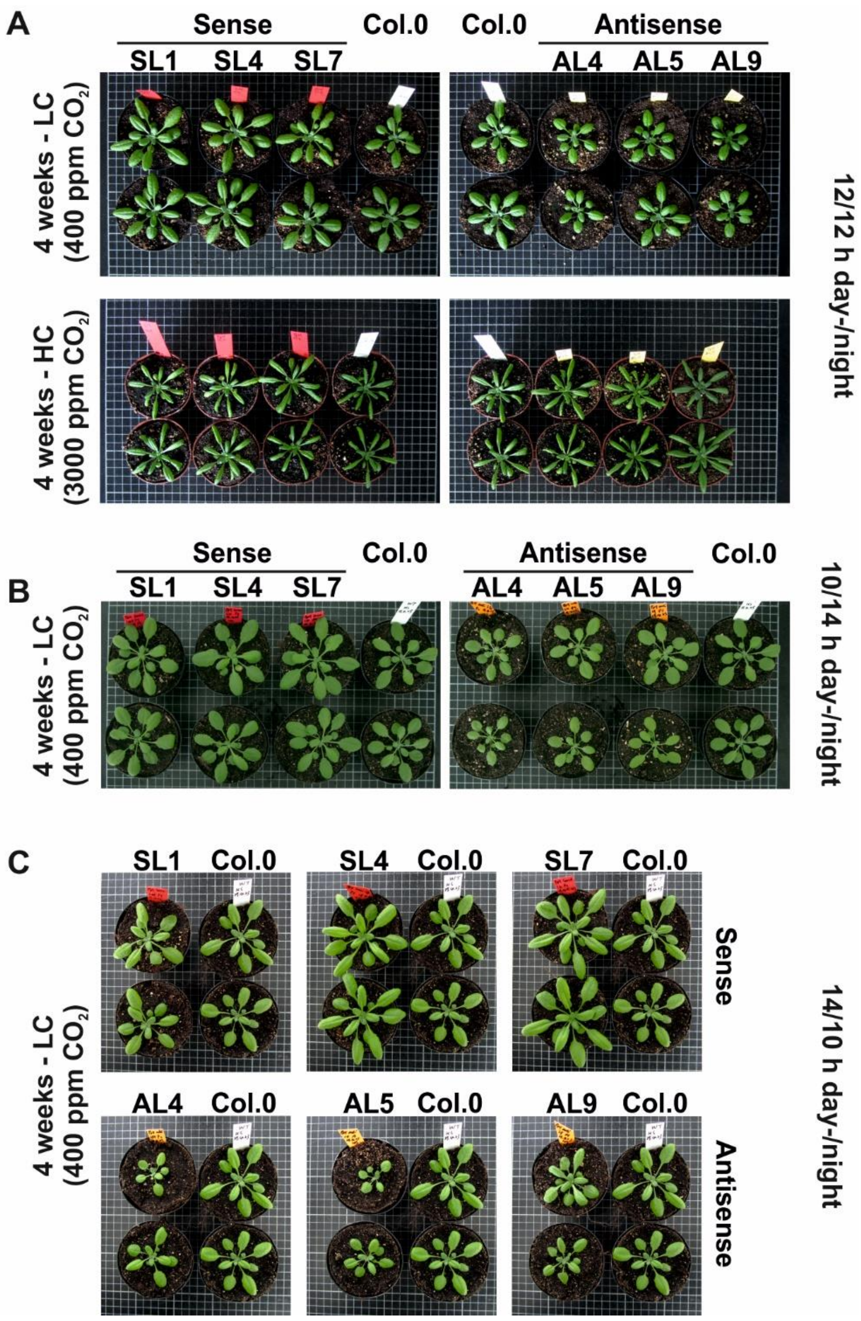


**Supplemental Figure S2.** Phenotype of Arabidopsis GC specific GDC-H modulated lines and the wildtype under different growth conditions.

**(A)** Representative images of plants grown for 4 weeks in normal air (upper panel: LC – low carbon; 400 ppm $CO_2$) or in elevated $CO_2$ (lower panel: HC – low carbon; 3000 ppm $CO_2$) in a 12/12 h day-/night-cycle. Representative images of plants grown for 4 weeks in normal air (LC – low carbon; 400 ppm $CO_2$) in a **(B)** 10/14 h and **(C)** 14/10 h day-/night-cycle with otherwise equal conditions.

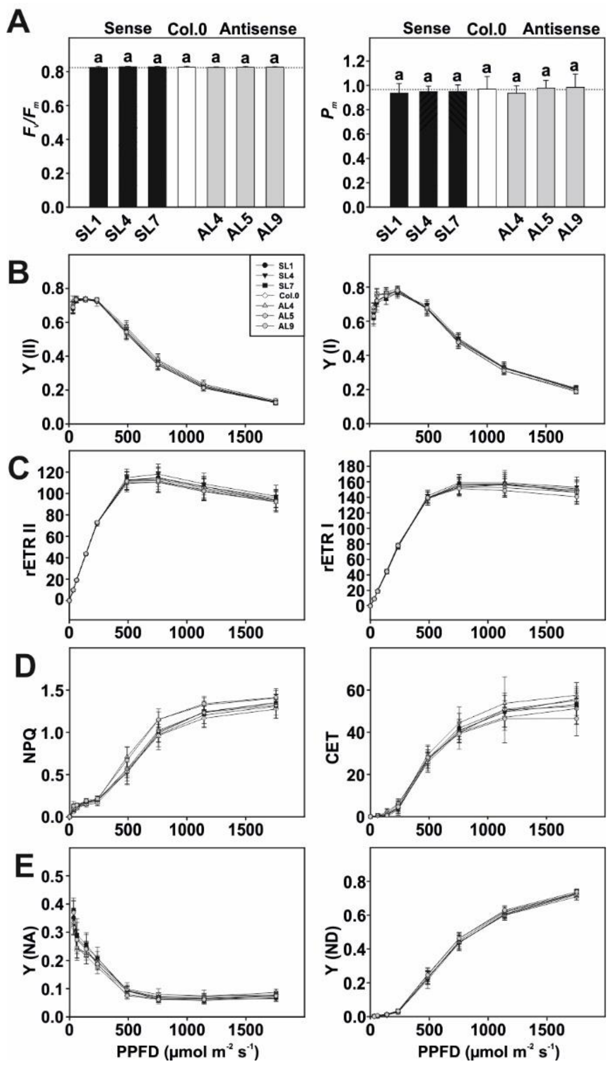


**Supplemental Figure S3.** Chlorophyll fluorescence parameters of Arabidopsis GC specific GDC-H modulated lines and the wildtype.

Displayed are selected parameters associated with PSII and PSI integrity and functioning of all genotypes grown under standard growth conditions to stage 5.1 (Boyes et al., 2001). Given are: **(A)** Maximum efficiency of PSII ($F_v/F_m$) and maximum oxidizable P700 ($P_m$) from dark adapted plants; **(B)** photosynthetic efficiency curves of PSII (Y[II]) and PSI (Y[I]); **(C)** relative electron transport rates of PSII (rETR II) and PSI (rETR I); **(D)** non-photochemical quenching of PSII (NPQ) and cyclic electron flow around PSI (CET); and, **(E)** acceptor (Y [NA]) and donor (Y [ND]) side limitation of PSI. Shown are means ± SD of at least 6 biological replicates. Values that do not share the same letter are significantly different from each other as determined by ANOVA. Note, lack of letters in B to E is explained due to absence of statistical differences.

### Supplemental Tables

**Supplemental Table S1.** Light response curves of the transgenic lines and the wildtype under standard conditions.

Plants were grown under environmental controlled conditions in normal air (400 ppm $CO_2$) to growth stage 5.1 (Boyes et al., 2001). Light response curves were measured from high-to-low light (1759,1144,757,488,236,143,62,36, and 0 µmol $m^{-2}$ $s^{-1}$) following 10 min of light adaptation (1000 µmol $m^{-2}$ $s^{-1}$). Given are: net $CO_2$ assimilation rate ($A_N$), stomatal conductance ($g_s$), intracellular $CO_2$ concentration ($C_i$), transpiration rate ($E$); and intrinsic water use efficiency (WUE). Values are means ± SD (n = 6). Values that do not share the same letter are significantly different from each other as determined by ANOVA.

| | | Light intensity (µmol $m^{-2}$ $s^{-1}$) | | | | | | | | |
|---|---|---|---|---|---|---|---|---|---|---|
| | Line | 0 | 36 | 62 | 143 | 236 | 488 | 757 | 1144 | 1759 |
| | SL1 | - 0.15 ± 0.60[a] | 1.87 ± 0.48[a] | 3.02 ± 0.53[a] | 5.63 ± 6.98[a] | 6.99 ± 0.86[a] | 8.11 ± 0.76[b] | 8.39 ± 0.69[b] | 8.65 ± 0.73[b] | 8.88 ± 0.40[b] |
| | SL4 | - 0.25 ± 0.91[a] | 1.76 ± 1.11[a] | 2.59 ± 1.34[ab] | 5.64 ± 1.36[a] | 7.15 ± 1.08[a] | 8.48 ± 1.09[b] | 8.76 ± 1.00[b] | 8.84 ± 1.05[b] | 9.08 ± 0.89[b] |
| $A_N$ | SL7 | - 0.1 ± 1.14[a] | 2.25 ±0.99[a] | 2.95 ± 1.12[ab] | 6.07 ± 0.75[a] | 7.40 ± 1.29[a] | 8.57 ± 0.96[b] | 9.03 ± 0.80[b] | 9.33 ± 0.82[b] | 9.19 ± 1.03[b] |
| (µmol $CO_2$ $m^{-2}$ $s^{-1}$) | Col.0 | - 0.16 ± 0.74[a] | 1.82 ± 0.79[a] | 2.62 ± 0.86[ab] | 5.20 ± 0.29[a] | 6.45 ± 0.44[a] | 7.05 ± 0.61[a] | 7.70 ± 0.63[a] | 7.92 ± 0.25[a] | 7.90 ± 0.44[a] |
| | AL4 | - 0.32 ± 0.49[a] | 1.03 ± 0.25[a] | 2.27 ± 0.51[b] | 4.31 ± 0.34[b] | 5.36 ± 0.50[b] | 5.98 ± 0.51[c] | 6.09 ± 0.73[c] | 6.36 ± 0.36[c] | 6.41 ± 0.34[c] |
| | AL5 | - 0.80 ± 0.44[a] | 1.27 ± 0.16[a] | 2.32 ± 0.16[b] | 4.93 ± 0.69[ab] | 5.88 ± 0.69[b] | 6.57 ± 0.43[c] | 6.69 ± 0.45[c] | 7.16 ± 0.77[c] | 7.15 ± 0.51[c] |
| | AL9 | - 0.83 ± 0.44[a] | 1.68 ± 0.31[a] | 1.98 ± 0.32[b] | 4.49 ± 0.64[b] | 5.53 ± 0.41[b] | 6.35 ± 0.43[c] | 6.42 ± 0.93[c] | 6.60 ± 0.36[c] | 6.80 ± 0.37[c] |
| | SL1 | 0.17 ± 0.07[a] | 0.20 ± 0.07[a] | 0.23 ± 0.07[a] | 0.26 ± 0.06[b] | 0.30 ± 0.05[b] | 0.33 ± 0.05[b] | 0.36 ± 0.05[b] | 0.38 ± 0.05[b] | 0.40 ± 0.05[b] |
| | SL4 | 0.18 ± 0.04[a] | 0.21 ± 0.04[a] | 0.24 ± 0.04[a] | 0.28 ± 0.04[b] | 0.30± 0.03[b] | 0.33 ± 0.03[b] | 0.35 ± 0.03[b] | 0.37 ± 0.03[b] | 0.39 ± 0.03[b] |
| $g_s$ | SL7 | 0.17 ± 0.03[a] | 0.19 ± 0.04[a] | 0.23 ± 0.04[a] | 0.26 ± 0.04[b] | 0.28 ± 0.04[b] | 0.31 ± 0.04[b] | 0.33 ± 0.04[b] | 0.35 ± 0.04[b] | 0.37 ± 0.03[b] |
| (mol $m^{-2}$ $s^{-1}$) | Col.0 | 0.14 ± 0.03[a] | 0.16 ± 0.03[a] | 0.19 ± 0.03[a] | 0.20 ± 0.03[a] | 0.22 ± 0.03[a] | 0.23 ± 0.03[a] | 0.25 ± 0.03[a] | 0.26 ± 0.02[a] | 0.28 ± 0.03[a] |
| | AL4 | 0.08 ± 0.03[b] | 0.09 ± 0.03[b] | 0.11 ± 0.04[b] | 0.13 ± 0.04[c] | 0.14 ± 0.04[c] | 0.16 ± 0.04[c] | 0.17 ± 0.04[c] | 0.19 ± 0.04[c] | 0.21 ± 0.03[c] |
| | AL5 | 0.08 ± 0.02[b] | 0.10 ± 0.02[b] | 0.12 ± 0.03[b] | 0.13 ± 0.03[c] | 0.15 ± 0.03[c] | 0.16 ± 0.03[c] | 0.18 ± 0.03[c] | 0.20 ± 0.02[c] | 0.22 ± 0.02[c] |
| | AL9 | 0.06 ± 0.02[b] | 0.07 ± 0.02[b] | 0.08 ± 0.03[b] | 0.10 ± 0.03[c] | 0.11 ± 0.03[c] | 0.12 ± 0.04[c] | 0.14 ± 0.04[c] | 0.16 ± 0.04[c] | 0.19 ± 0.04[c] |
| | SL1 | 394.1 ± 8.1[a] | 378.3 ± 7.3[b] | 371.4 ± 3.6[b] | 355.3± 4.7[b] | 351.4 ± 4.5[b] | 345.6 ± 6.0[b] | 346.6 ± 5.4[b] | 346.9 ± 4.0[b] | 346.9 ± 3.3[b] |

| | | | | | | | | | | |
|---|---|---|---|---|---|---|---|---|---|---|
| | SL4 | 392.6 ± 8.1$^{a}$ | 381.1 ± 8.9$^{b}$ | 376.7 ± 6.04$^{b}$ | 358.3 ± 4.5$^{b}$ | 351.3 ± 3.6$^{b}$ | 346.2 ± 2.7$^{b}$ | 346.4± 2.1$^{b}$ | 347.6 ± 3.0$^{b}$ | 347.4 ± 1.9$^{b}$ |
| $C_i$ | SL7 | 390.5 ± 12.6$^{a}$ | 377.6 ± 10.7$^{ab}$ | 375.2 ± 3.29$^{b}$ | 353.1 ± 1.0$^{b}$ | 350.1 ± 4.5$^{b}$ | 344.3 ± 2.5$^{b}$ | 342.6± 3.1$^{b}$ | 342.6 ± 1.2$^{b}$ | 344.6± 2.4$^{ab}$ |
| (μmol) | Col.0 | 393.1 ± 5.5$^{a}$ | 372.3 ± 1.9$^{a}$ | 367.0 ± 5.8$^{a}$ | 346.2 ± 8.6$^{a}$ | 339.1 ± 8.7$^{a}$ | 337.6 ± 4.8$^{a}$ | 335.8 ± 6.7$^{a}$ | 336.6 ± 3.7$^{a}$ | 338.8 ± 6.7$^{a}$ |
| | AL4 | 397.1 ± 12.4$^{a}$ | 370.5 ± 9.1$^{a}$ | 345.4 ± 12.2$^{c}$ | 331.0 ± 12.6$^{a}$ | 324.2± 13.5$^{c}$ | 323.8 ± 9.6$^{c}$ | 329.5 ± 9.3$^{c}$ | 332.1 ± 8.7$^{ac}$ | 336.7 ± 8.6$^{a}$ |
| | AL5 | 408.6 ± 7.8$^{a}$ | 367.9 ± 8.9$^{a}$ | 354.5 ± 14.2$^{c}$ | 325.2 ± 17.9$^{c}$ | 320.9 ± 17.4$^{c}$ | 320.0 ± 13.6$^{c}$ | 325.2 ± 10.2$^{c}$ | 325.6 ± 11.1$^{c}$ | 331.4 ± 8.1$^{ac}$ |
| | AL9 | 417.1± 13.9$^{a}$ | 344.3 ± 20.2$^{c}$ | 344.6 ± 25.4$^{c}$ | 305.7 ± 20.5$^{c}$ | 298.9 ± 34.1$^{c}$ | 294.7 ±31.5$^{c}$ | 306.5± 24.0$^{c}$ | 318.7 ± 20.5$^{c}$ | 324.6 ± 14.2$^{c}$ |
| | SL1 | 2.24 ± 0.56$^{a}$ | 2.54 ± 0.55$^{a}$ | 2.88 ± 0.52$^{a}$ | 3.28 ± 0.53$^{a}$ | 3.69 ± 0.57$^{b}$ | 4.15 ± 0.60$^{b}$ | 4.58 ± 0.63$^{b}$ | 5.05 ± 0.64$^{b}$ | 5.65 ± 0.65$^{b}$ |
| | SL4 | 2.30 ± 0.43$^{a}$ | 2.59 ± 0.38$^{a}$ | 2.93 ± 0.34$^{a}$ | 3.29 ± 0.32$^{a}$ | 3.61 ± 0.31$^{b}$ | 4.00 ± 0.30$^{b}$ | 4.31 ± 0.31$^{b}$ | 4.73 ± 0.33$^{b}$ | 5.18 ± 0.29$^{b}$ |
| $E$ | SL7 | 2.07 ± 0.31$^{a}$ | 2.33 ± 0.33$^{a}$ | 2.64 ± 0.34$^{a}$ | 2.97 ± 0.34$^{a}$ | 3.27 ± 0.34$^{b}$ | 3.64 ± 0.32$^{b}$ | 3.98 ± 0.33$^{b}$ | 4.43 ± 0.39$^{b}$ | 4.83 ± 0.33$^{b}$ |
| (μmol $H_2O$ $m^{-2}$ $s^{-1}$) | Col.0 | 1.80 ± 0.39$^{a}$ | 2.01 ± 0.41$^{a}$ | 2.25 ± 0.37$^{a}$ | 2.47 ± 0.39$^{a}$ | 2.67 ± 0.38$^{a}$ | 2.92 ± 0.35$^{a}$ | 3.21 ± 0.35$^{a}$ | 3.52 ± 0.35$^{a}$ | 3.96 ± 0.40$^{a}$ |
| | AL4 | 1.01 ± 0.38$^{b}$ | 1.17 ± 0.43$^{b}$ | 1.41 ± 0.46$^{b}$ | 1.61 ± 0.49$^{b}$ | 1.82 ± 0.48$^{c}$ | 2.08 ± 0.51$^{c}$ | 2.36 ± 0.53$^{c}$ | 2.68 ± 0.55$^{c}$ | 3.05 ± 0.53$^{c}$ |
| | AL5 | 1.00 ± 0.27$^{b}$ | 1.19 ± 0.29$^{b}$ | 1.42 ± 0.34$^{b}$ | 1.66 ± 0.34$^{b}$ | 1.87 ± 0.34$^{c}$ | 2.12 ± 0.33$^{c}$ | 2.41 ± 0.30$^{c}$ | 2.76 ± 0.23$^{c}$ | 3.16 ± 0.19$^{c}$ |
| | AL9 | 0.68 ± 0.25$^{b}$ | 0.81 ± 0.26$^{b}$ | 1.01 ± 0.34b | 1.18 ± 0.37$^{b}$ | 1.39 ± 0.41$^{c}$ | 1.58 ± 0.50$^{c}$ | 1.86 ± 0.56$^{c}$ | 2.20 ± 0.55$^{c}$ | 2.66 ± 0.55$^{c}$ |
| | SL1 | - 0.19 ± 0.39$^{a}$ | 0.82 ± 0.26$^{a}$ | 1.11 ± 0.22$^{a}$ | 1.80 ± 0.29$^{a}$ | 1.97 ± 0.27$^{b}$ | 2.01 ± 0.27$^{b}$ | 1.88 ± 0.23$^{b}$ | 1.75 ± 0.20$^{b}$ | 1.61 ± 0.20$^{b}$ |
| | SL4 | - 0.01 ± 0.47$^{a}$ | 0.77 ± 0.51$^{a}$ | 0.98 ± 0.52$^{a}$ | 1.81 ± 0.48$^{a}$ | 2.06 ± 0.42$^{ab}$ | 2.20 ± 0.36$^{ab}$ | 2.08 ± 0.32$^{ab}$ | 1.92 ± 0.30$^{ab}$ | 1.80 ± 0.23$^{ab}$ |
| *WUE* | SL7 | - 0.16 ± 0.58$^{a}$ | 1.04 ± 0.54$^{a}$ | 1.20 ± 0.52$^{a}$ | 2.09 ± 0.41$^{a}$ | 2.30 ± 0.50$^{ab}$ | 2.41 ± 0.35$^{ab}$ | 2.31 ± 0.28$^{a}$ | 2.14 ± 0.24$^{a}$ | 1.94 ± 0.25$^{ab}$ |
| (μmol $CO_2$ $mol^{-1}$ $H_2O$) | Col.0 | - 0.03 ± 0.43$^{a}$ | 0.95 ± 0.40$^{a}$ | 1.17 ± 0.39$^{a}$ | 2.22 ± 0.49$^{ab}$ | 2.54 ± 0.47$^{ac}$ | 2.49 ± 0.30$^{ac}$ | 2.51 ± 0.40$^{a}$ | 2.31 ± 0.27$^{a}$ | 2.07 ± 0.35$^{ac}$ |
| | AL4 | - 0.41 ± 0.53$^{a}$ | 0.98 ± 0.40$^{a}$ | 1.59 ± 0.49$^{a}$ | 2.73 ± 0.54$^{b}$ | 3.00 ± 0.56$^{cd}$ | 2.95 ± 0.42$^{c}$ | 2.61 ± 0.38$^{ac}$ | 2.45 ± 0.39$^{ac}$ | 2.14 ± 0.36$^{ac}$ |
| | AL5 | - 0.80 ± 0.42$^{a}$ | 1.10 ± 0.39$^{a}$ | 1.72 ± 0.60$^{a}$ | 3.01 ± 0.73$^{c}$ | 3.21 ± 0.69$^{d}$ | 3.19 ± 0.52$^{cd}$ | 2.84 ± 0.38$^{cd}$ | 2.63 ± 0.38$^{c}$ | 2.30 ± 0.19$^{cd}$ |
| | AL9 | - 1.30 ± 0.58$^{a}$ | 1.95 ± 0.82$^{a}$ | 1.97 ± 0.86$^{a}$ | 3.60 ± 0.75$^{c}$ | 3.95 ± 0.98$^{d}$ | 4.10 ± 1.04$^{d}$ | 3.37 ± 0.44$^{d}$ | 2.80 ± 0.40$^{c}$ | 2.61 ± 0.44$^{d}$ |

**Supplemental Table S2.** Calculated parameters from light response curves numerically given in Suppplemental Table S1.

Estimations of the maximum photosynthetic rate ($A_{max}$) and initial slopes of the light response curves ($\alpha_p$) from the light response curves measured from the transgenic lines in comparison with the wildtype. $A_{max}$ showed ~10-15% increases in overexpression and ~15-19% decreases in antisense lines. Values of $\alpha_p$ were accelerated to about 40% in overexpressors, but remained significantly unchanged in the antisense suppressors compared to the wildtype. Given are means ± SD (n = 6). Values that do not share the same letter are significantly different from each other as determined by ANOVA.

| | Parameter | |
|---|---|---|
| Genotype | $A_{max}$ | $\alpha_p$ |
| SL1 | **8.98 ± 0.87[b]** | **0.14 ± 0.05[b]** |
| SL4 | **9.26 ± 0.86[b]** | **0.14 ± 0.01[b]** |
| SL7 | **9.40 ± 1.07[b]** | **0.13 ± 0.02[b]** |
| Col.0 | 8.15 ± 0.43[a] | 0.09 ± 0.01[a] |
| AL4 | **6.57 ± 0.64[c]** | 0.08 ± 0.01[a] |
| AL5 | **7.05 ± 0.56[c]** | 0.08 ± 0.01[a] |
| AL5 | **6.94 ± 0.39[c]** | 0.09 ± 0.01[a] |

**Supplemental Table S3.** Abundances of selected intermediates associated with primary metabolism in the transgenic lines and the wildtype under standard conditions.

Plants were grown under environmental controlled conditions in normal air (400 ppm $CO_2$) to growth stage 5.1 (Boyes et al., 2001). Leaf-material was harvested at the end of the day (11 h illumination) and frozen in liquid nitrogen until LC-MS/MS analysis. Given are absolute contents (nmol * mg $DW^{-1}$) of 35 primary metabolites and the sum parameters of the total amino acid (AAs) and organic acid (OAs) amounts (both in µmol * mg $DW^{-1}$). Values are mean ± SD from 6-8 biological replicates. Values that do not share the same letter are significantly different from each other as determined by ANOVA.

| | Genotype | | | | | | |
|---|---|---|---|---|---|---|---|
| **Metabolite** | **SL1** | **SL4** | **SL7** | **Col.0** | **AL4** | **AL5** | **AL9** |
| 2PG | **196.81 ±14.20[b]** | **200.48 ± 16.75[b]** | **199.09 ± 4.81[b]** | 222.61 ±7.1[a] | 236.6 ±19.77[a] | 250.39 ± 9.14[a] | 244.86 ± 22.09[a] |
| 3PGA | **86.94 ± 7.19[b]** | **82.71 ± 77.32[b]** | **77.32 ± 6.04[b]** | 54.58 ± 15.744[a] | **38.46 ± 7.86[c]** | **29.70 ± 17.10[c]** | 41.15 ± 12.87[ac] |
| Aconitate | 2754.10 ± 713.83[a] | 3210.59 ± 276.55[a] | 3493.71 ± 393.75[a] | 3613.72 ± 902.13[ac] | **4453.99 ± 590.05[c]** | **4573.96 ± 700.63[c]** | **4220.98 ± 915.78[c]** |
| Alanine | **4931.78 ± 129.84[b]** | 5446.78 ± 773.79[ab] | 6124.59 ± 322.22[a] | 5757.27 ± 546.31[a] | **7095.14 ± 453.57[c]** | **7589.52 ± 597.03[c]** | **7186.99 ± 1257.59[c]** |
| AMP | 2178.56 ± 809.27[a] | 1734.05 ±269.6[a] | 1954.39 ± 498.04[a] | 1424.01 ± 408.69[a] | 1568.99 ± 387.52[a] | 1980.35 ± 387.52[a] | 1851.10 ± 341.53[a] |
| Arginine | 1358.01 ± 367.92[a] | 1440.47 ± 307.67[a] | 1594.13 ± 450.36[a] | 1698.54 ± 419.34[a] | 1478.54 ± 262.34[a] | 1288.99 ± 383.93[a] | 1160.71 ± 267.49[a] |
| Asparagine | 4143.71 ± 430.32[a] | 4816.02 ± 927.75[a] | 4922.18 ± 457.53[a] | 5491.46 ± 979.22[a] | 4508.90± 430.88[a] | 4062.29 ± 1168.40[a] | 4000.45 ± 779.43[a] |
| Citrate | **9710.44 ± 1966.83[b]** | **10140.88 ± 881.71[b]** | **11050.46 ± 869.62[b]** | 13179.15 ± 566.06[a] | **16943.86 ± 1741.22[c]** | **17274.68 ± 2182.32[c]** | **17782.02 ± 2924.88[c]** |
| Citrulline | 56894.57 ± 13109.11[a] | 47330.12 ± 5142.14[a] | 46530.39 ± 8804.66[a] | 41437.25 ± 5318.67[a] | 45094.53 ± 3425.93[a] | 45283.32 ± 9490.64[a] | 42009.70 ± 3305.33[a] |
| Cysteine | 5808.76 ± 1210.64[a] | 6169.45 ± 702.78[a] | 7444.71 ± 1812.34[a] | 6707.67 ± 1185.68[a] | 7138.04 ± 1098.89[a] | 6389.58 ± 598.47[a] | 6427.24 ± 778.43[a] |
| Cystine | **23.25 ±5.65[b]** | **20.46 ± 5.41[b]** | **18.76 ± 4.44[b]** | 15.75 ± 3.20[a] | 15.28 ± 3.58[a] | **11.27 ± 2.46[c]** | **10.84 ± 1.88[c]** |
| Fumarate | **1507.69 ± 310.44[b]** | 2140.43 ±395.32[ab] | **1780.82 ± 461.14[b]** | 2429.35 ± 379.06[a] | **1897.18 ± 313.35[b]** | **1920.82 ± 301.28[b]** | **1682.45 ± 476.31[b]** |
| GABA | 225.32 ± 97.69[a] | 274.87 ± 87.89[a] | 377.56 ± 96.13[a] | 358.52 ± 128.29[a] | 347.33 ± 214.81[a] | 337.59 ± 200.61[a] | 260.92 ± 97.29[a] |

| | | | | | | | |
|---|---|---|---|---|---|---|---|
| Glutamate | 56894.59 ± 13109.11$^{a}$ | 47330.12 ± 5142.14$^{a}$ | 46530.39 ± 8804.66$^{a}$ | 41437.25 ± 5318.67$^{a}$ | 45094.53 ± 3425.92$^{a}$ | 45283.32 ± 9490.64$^{a}$ | 42009.70 ± 3305.33$^{a}$ |
| Glutamine | 44557.82 ± 6912.19$^{a}$ | 40982.37 ± 3583.99$^{a}$ | 45748.10 ± 4260.41$^{a}$ | 49502.05 ± 8322.49$^{a}$ | 46542.89 ± 1939.36$^{a}$ | 45381.72 ± 7699.53$^{a}$ | 38783.80± 5182.57$^{a}$ |
| Glycine | 1475.93 ± 252.54$^{a}$ | 1365.98 ± 193.24$^{a}$ | 1338.77 ± 129.42$^{a}$ | 1413.01 ± 268.00$^{a}$ | 1408.51 ± 186.99$^{a}$ | 1453.26 ± 305.43$^{a}$ | 1387.74 ± 329.99$^{a}$ |
| Histidine | 265.11 ± 17.12$^{a}$ | 312.46 ± 34.93$^{a}$ | 317.85 ± 21.30$^{a}$ | 355.90 ± 47.10$^{a}$ | 316.26 ± 49.13$^{a}$ | 323.66 ± 71.21$^{a}$ | 314.79 ± 56.50$^{a}$ |
| Isocitrate | **9359.07 ± 1011.51$^{b}$** | **10306.22 ± 951.74$^{b}$** | **11540.69 ± 861.15$^{b}$** | 12761.97 ± 1268.35$^{a}$ | **15920.15 ± 1546.35$^{c}$** | **15968.42 ± 2118.45$^{c}$** | **16666.64 ± 2192.15$^{c}$** |
| Isoleucine | 319.84 ± 61.51$^{a}$ | 362.26 ± 69.26$^{a}$ | 368.26± 82.57$^{a}$ | 384.66 ± 59.91$^{a}$ | 325.96 ± 47.74$^{a}$ | 346.57 ± 95.55$^{a}$ | 328.82 ± 67.08$^{a}$ |
| Lactate | **10112.69 ± 1104.84$^{b}$** | **10319.02 ± 744.94$^{b}$** | **11322.86 ± 876.12$^{b}$** | 13725.08 ± 1385.62$^{a}$ | **17645.74 ± 1813.35$^{c}$** | **17990.27 ± 2272.72$^{c}$** | **18518.62 ± 2600.99$^{c}$** |
| L-Arg succinic acid | **22.15 ± 5.83$^{b}$** | **19.53 ± 2.69$^{b}$** | **20.04 ± 1.83$^{b}$** | 29.88 ± 3.94$^{a}$ | **20.49 ± 3.66$^{b}$** | **18.25 ± 3.20$^{b}$** | **16.4 ± 1.65$^{b}$** |
| Leucine | 306.68 ± 59.44$^{a}$ | 384.13 ± 76.44$^{a}$ | 362.84 ± 84.50$^{a}$ | 375.44 ± 82.09$^{a}$ | 306.58 ± 58.77$^{a}$ | 346.92 ± 99.10$^{a}$ | 309.64 ± 76.42$^{a}$ |
| Lysine | **51030.16 ± 8602.32$^{b}$** | 46903.89 ± 4523.16$^{a}$ | 52501.26 ± 4870.72$^{a}$ | 56406.97 ± 9179.43$^{a}$ | 53142.93 ± 2248.53$^{a}$ | 48076.16 ± 13981.99$^{a}$ | 44722.09 ± 5803.52$^{a}$ |
| Malate | **100890.11 ± 4794.20$^{b}$** | **128883.34 ± 12481.41$^{b}$** | **117954.73 ± 6698.90$^{b}$** | 154926.93 ± 5832.78$^{a}$ | 153800.31 ± 5888.59$^{a}$ | **170983.67 ± 4788.51$^{c}$** | **175962.43 ± 11303.01$^{c}$** |
| Methionine | 550.50 ± 116.19$^{a}$ | 448.87 ± 76.58$^{a}$ | 464.89 ± 50.34$^{a}$ | 517.14 ± 100.66$^{a}$ | 576.61 ± 63.52$^{a}$ | 508.67 ± 161.24$^{a}$ | 522.09 ± 96.01$^{a}$ |
| NAD | 246.22 ± 22.06$^{a}$ | 246.24 ± 34.41$^{a}$ | 209.23 ± 50.31$^{a}$ | 287.86 ± 44.85$^{ab}$ | 344.29 ± 60.32$^{b}$ | 346.13 ±50.24$^{b}$ | 299.53 ± 30.22$^{b}$ |
| Ornithine | 174.23 ± 27.28$^{a}$ | 169.76 ± 35.83$^{a}$ | 167.06 ± 17.88$^{a}$ | 185.84 ± 40.86$^{a}$ | 151.88 ± 15.16$^{a}$ | 133.07 ± 39.49$^{a}$ | 127.78 ± 23.47$^{a}$ |
| Phenylalanine | 1920.20 ± 408.50$^{a}$ | 1987.99 ± 383.68$^{a}$ | 2155.27 ± 502.93$^{a}$ | 2011.63 ± 258.12$^{a}$ | 1804.22 ± 306.66$^{a}$ | 2011.00 ± 542.23$^{a}$ | 1863.67 ± 482.59$^{a}$ |
| Proline | 7404.10 ± 4465.44$^{ab}$ | 13730.77± 6990.52$^{ab}$ | 9183.13 ± 5580.81$^{ab}$ | 16081.86 ± 6754.23$^{a}$ | 12872.53 ± 6216.91$^{ab}$ | **6058.79 ± 2383.81$^{b}$** | **6442.07 ± 3062.62$^{b}$** |
| Serine | 10146.54 ± 1601.52$^{a}$ | 9951.51 ± 1691.72$^{a}$ | 9912.75 ± 1276.59$^{a}$ | 10087.77 ± 633.51$^{a}$ | 9541.70 ± 1794.58$^{a}$ | 9673.16 ± 1331.07$^{a}$ | 10443.10 ± 2360.14$^{a}$ |
| Succinate | **200.64 ± 24.14$^{b}$** | **305.26 ± 41.96$^{c}$** | **270.45 ± 70.27$^{bc}$** | 404.65 ± 63.11$^{a}$ | **288.22 ± 38.79$^{bc}$** | **293.16 ± 71.70$^{bc}$** | **263.26 ± 87.43$^{bc}$** |
| Threonine | 6986.76 ± 954.96$^{a}$ | 6550.99 ± 585.58$^{a}$ | 7383.66 ± 1128.46$^{a}$ | 6695.67 ± 1021.55$^{a}$ | 7127.99 ± 379.77$^{a}$ | 7312.22 ± 1138.18$^{a}$ | 7036.77 ± 870.30$^{a}$ |
| Tryptophan | 252.19 ± 42.58$^{a}$ | 305.25 ± 51.50$^{a}$ | 273.32± 23.17$^{a}$ | 307.33 ± 64.98$^{a}$ | 237.88 ± 39.47$^{a}$ | 252.63 ± 79.44$^{a}$ | 254.38 ± 42.45$^{a}$ |
| Tyrosine | **115.82 ± 18.52$^{b}$** | **118.51 ± 16.54$^{b}$** | **119.75 ± 6.72$^{b}$** | 96.84 ± 14.75$^{a}$ | 86.78 ± 22.09$^{ac}$ | **84.11 ± 15.22$^{c}$** | 93.03 ± 12.81$^{ac}$ |
| Valine | 225.32 ± 97.69$^{a}$ | 274.87 ± 87.89$^{a}$ | 377.56 ± 96.13$^{a}$ | 358.52 ± 128.29$^{a}$ | 347.34 ± 214.81$^{a}$ | 337.59 ± 200.61$^{a}$ | 260.92 ± 97.29$^{a}$ |
| Total AAs | **271513.3 ± 28580.4$^{b}$** | 254674.3 ± 17867.1$^{ab}$ | **251666.9 ± 11728.4$^{b}$** | 232065.5 ± 13637.7$^{a}$ | 231977.0 ± 14938.5$^{abc}$ | **212502.2 ± 13603.4$^{c}$** | **205638.8 ± 15105.6$^{c}$** |
| Total OAs | **129751.8 ± 9976.9$^{c}$** | **163534.8 ± 13250.7$^{b}$** | **160207.1 ± 14629.5$^{b}$** | 199029.1 ± 9697.6$^{a}$ | 194965.9 ± 13126.0$^{a}$ | 215609.6 ± 19762.4$^{a}$ | 215178.1 ± 46209.9$^{a}$ |

**Supplemental Table S4.** Loadings of metabolites on the first three principal components (PCs) in leaves of the GC specific GDC-H lines and the wildtype.

Summary of the numerical values of the loadings on the first three PCs of the PCA shown in Figure 4 (C and D) are given. Loadings with the absolute values > ± 0.2 (strong impact) are shown in bold.

| **Metabolite** | **PC1** | **PC2** | **PC3** |
|---|---|---|---|
| Total organic acids | **-0.44632** | 0 | 0 |
| Fructose | **0.36208** | 0 | 0 |
| Glucose | **0.33913** | 0 | 0 |
| Citrate | **-0.32666** | 0 | 0 |
| Lactate | **-0.32666** | 0 | 0 |
| Isocitrate | **-0.31159** | 0 | 0 |
| Sucrose | **0.28589** | 0 | 0 |
| Total amino acids | **0.27658** | 0 | 0 |
| Starch | **0.27106** | 0 | 0 |
| Malate | -0.11172 | 0 | -0.043679 |
| Succinate | 0 | **-0.58325** | 0 |
| Aconitate | 0 | **-0.58325** | 0 |
| Proline | 0 | **-0.33807** | 0 |
| Fumarate | 0 | **-0.30599** | 0 |
| Asparagine | 0 | **-0.22023** | 0 |
| AMP | 0 | 0.17414 | 0 |
| Alanine | 0 | 0.13506 | 0 |
| Histidine | 0 | -0.098297 | 0 |
| Glutamate | 0 | 0.06054 | 0 |
| Citrulline | 0 | 0.03658 | 0 |
| Arginine | 0 | 0 | **0.26612** |
| Cysteine | 0 | 0 | **0.23254** |
| Cystine | 0 | 0 | 0.11401 |
| Glutamine | 0 | 0 | **0.73563** |
| Glycine | 0 | 0 | 0 |
| Isoleucine | 0 | 0 | 0 |
| Leucine | 0 | 0 | 0 |
| Lysine | 0 | 0 | **0.49949** |
| Methionine | 0 | 0 | 0 |
| Phenylalanine | 0 | 0 | 0 |
| Serine | 0 | 0 | -0.12492 |
| Threonine | 0 | 0 | 0 |
| Tryptophan | 0 | 0 | 0 |
| Tyrosine | 0 | 0 | -0.021008 |
| Valine | 0 | 0 | 0.030927 |

| | | | |
|---|---|---|---|
| GABA | 0 | 0 | 0 |
| 2-PG | 0 | 0 | 0 |
| 3-PGA | 0 | 0 | 0 |
| L-Argininosuccinic acid | 0 | 0 | **0.22923** |
| NAD | 0 | 0 | 0 |
| Ornithine | 0 | 0 | 0 |

**Supplemental Table S5.** Primers used for PCR amplification of genomic DNA and cDNA.

Underlined sequences indicate the introduced *BamHI*, *SacI* and *PstI* sites in the primers used to produce expression constructs. ATG in bold print highlight the start codon for methionine.

| Stock Number | Name | Sequence (5´-to3´) |
|---|---|---|
| P950 | *At*GC1_S1141_SacI | GAGCTC**ATG**GTTGCAACAGAGAGGATGAAT |
| P951 | *At*GC1-AS-BamHI | GGATCCATTTCTTGAGTAGTGATTTTGAAG |
| P965 | *Fp*GDCH-S-PstI | CTGCAG**ATG**GCTCTTAGAATCTGGGCT |
| P966 | *Fp*GDCH-AS-PstI | CTGCAGCTACGTGAGCAGAATCTTCTTC |
| P807 | 35STer | CTCGAGAGTATCGATCTGGATTTTAGT |
| P444 | *S16*-forward | GGCGACACAACCAGCTACTGA |
| P445 | *S16*-revers | CGGTAACTCTTCTGGTAACGA |
